# Mechanism-based enzyme activating compounds

**DOI:** 10.1101/2020.04.08.032235

**Authors:** Xiangying Guan, Alok Upadhyay, Raj Chakrabarti

## Abstract

Compared to inhibitors, which constitute the vast majority of today’s drugs, enzyme activators have considerable advantages, especially in the context of enzymes that regulate reactive flux through metabolic pathways associated with chronic, age-related diseases and lifespan. Across all families of enzymes, only a dozen or so distinct classes of small molecule activators have been characterized. Enzyme activators that are not based on naturally evolved allosteric mechanisms are much more difficult to design than inhibitors, because enzymatic catalysis has been optimized over billions of years of evolution. Here, we introduce modes of enzyme activation based on the catalytic reaction mechanisms of enzymes for which naturally evolved activators may not exist. We establish biophysical properties of small molecule modulators that are necessary to achieve desired changes in the steady state and non-steady state parameters of these enzymes, including changes in local conformational degrees of freedom conducive to the enhancement of catalytic activity that can be identified through computational modeling of their active sites. We illustrate how the modes of action of several compounds reported to activate enzymes without known allosteric sites may be understood using the framework presented. We also present simulations and new experimental results in support of this framework, including identification of the mechanism of a compound that activates the human SIRT3 enzyme, which does not contain a known allosteric site, under physiologically relevant conditions.

## I. INTRODUCTION

The vast majority of today’s drugs are enzyme inhibitors^1^. The mode of action of inhibitors generally involves transient knockdown of the function of a nonessential target enzyme in the context of treatment of a disease phenotype, whereas activators upregulate functions that are essential for a desired phenotype, such as enhanced healthspan or lifespan. Enzyme activators have considerable advantages over inhibitors particularly in geroscience research and drug development for age-related diseases^2,3^.

In order to determine when overexpression or activation of an enzyme will increase flux through the associated reactive pathway, methods from metabolic control theory^4^ (which include the control of biochemical reaction network dynamics^5^) can be used. Typically, over-expression or activation of an enzyme will not increase reactive pathway flux. Exceptions are so-called regulatory enzymes, whose overexpression or activation significantly increases the flux through the associated metabolic network. A small increase in activity of critical regulatory enzymes can have a dramatic, systemic effect of downstream pathways^6^ through a process of metabolic reprogramming. Such regulatory enzymes can play a critical role in age-related diseases and organismic longevity, as first demonstrated by studies in yeast^7^.The existence of naturally occurring allosteric activators reveals the functional significance of enzyme activation in the regulation of biochemical signaling pathways^2,3,8–11^. For example, the NAD+-dependent sirtuin deacylase SIRT1 and AMP-activated protein kinase both upregulate various transcriptional networks and contain allosteric binding sites^12^.

The enzymes AMP-activated protein kinase, nicotinamide phosphoribosyltransferase, and the mammalian sirtuins SIRT1-7 (silent information regulators 1-7) are core regulators of age-related diseases. Over the last several years, scientists have realized that the body regulates metabolism and lifespan in response to environmental cues through a tightly conserved axis consisting of these enzymes. This axis allows the body to react with stress defenses when nutrient availability is low (e.g., under caloric restriction), extending healthspan and lifespan through systemic changes that render energy utilization more efficient. Early studies showed the yeast sirtuin Sir2^7^ and mammalian SIRT1^3^ overexpression increase lifespan. It is now known that other sirtuins – including SIRT2, SIRT3 and SIRT6 –and multiple protein substrates play significant roles in regulating mammalian longevity^13–15^. The physiological importance of sirtuins has stimulated intense interest in designing sirtuin-activating compounds, several of which have been demonstrated to induce lifespan extension in model organisms such as mice^16,17^. In this regard the majority of work has been devoted to activators that allosterically target SIRT1. These compounds bind outside of the active site to an allosteric domain in SIRT1 that is not shared by SIRT2-7^12^. Moreover, allosteric activators only work with a limited set of SIRT1 substrates^3^.

Despite the historical focus on SIRT1 activators, several compounds have been reported in the literature as being activators of sirtuins other than SIRT1^18–20^. One such compound^20^ was also demonstrated to have physiologically rejuvenating effects consistent with the known effect of SIRT3 overexpression, reversing cardiac disease in mice. However, the modes of activation by these compounds are not fully understood, given that until recently^21^, no framework existed for small molecule modulation of this type.

Whereas theories of allosteric activation, such as the Monod Wyman Changeux (MWC) model, have been available since the 1960’s^22–24^, foundations for the rational design of non-allosteric, or *mechanism-based* enzyme activators have been lacking. Mechanism-based activation refers to a mode of enzyme activation wherein the modulator induces changes in local active site degrees of freedom, affecting multiple reaction steps. The kinetic effects of such compounds depend on the relative stabilities of the complexes of the activator with multiple reaction intermediates, necessitating the use of a complete steady-state or dynamic model for enzymatic catalysis to predict these effects. By contrast, allosteric activators function by inducing global conformational changes that positively affect one step of the reaction (e.g., reduction of substrate *K*_*d*_ or increase in chemical reaction rate). The kinetic effects of allosteric activators can be predicted without a complete steady state or dynamic model for catalysis. Allosteric activators typically rely on a naturally evolved mode of activation, whereas mechanism-based activation is a molecular engineering problem more akin to enzyme design^25^.

In this paper, we provide a full theoretical framework for the modes of action of mechanism-based enzyme activating compounds, as well as new experimental and simulation data in support of this theory. This framework extends the conventional classification of small molecule modulators of enzymes to encompass compounds that activate enzymes without binding to allosteric sites. We delineate differences between the biophysical modes of binding and active site modulation by this class of molecules and allosteric activators. This constitutes the first comprehensive framework available for characterization of mechanism-based enzyme activating compounds and as such is essential for development of such compounds into drugs, just as conventional inhibition theory is a foundation for standard drug development. One aspect of this theory, namely the kinetics of mechanism-based enzyme activation, was first introduced in approximate form without complete derivation in a recent paper from our group^21^. This paper goes further in several respects, both in terms of rigorous derivation of the kinetic theory of mechanism-based enzyme activation and the introduction of an atomistic free energy modulation theory of mechanism-based activation.

The paper is organized as follows. In Section 2, we describe how small molecules can induce modulation of local active site conformational degrees of freedom (as opposed to the global conformational changes in ensemble allosteric theory) to exert the effects on an enzyme’s catalytic free energy differences that are responsible for mechanism-based activation. In Section 3, we develop a kinetic formalism to delineate properties of mechanism-based enzyme activators for one and two substrate enzymes. In Section 4, we apply the kinetic theory of mechanism-based activation to an example enzyme (human SIRT3) for which non-allosteric activators have been reported and present new experimental data on one such hit compound (honokiol), identifying the mechanism whereby this compound activates this enzyme under physiologically relevant conditions and demonstrating that the experimental data are consistent with the activation framework presented herein. In Section 5, we present simulation results applying the structural theory of mechanism-based activation presented in Section 2 to the example enzyme.

## II. SMALL MOLECULE INDUCED MODULATION OF LOCAL PROTEIN DEGREES OF FREEDOM

The central differences between allosteric and mechanism-based enzyme activators, such as the fact that the probabilities of local rather than global conformational degrees of freedom are affected by modulator binding and the effects on steady state activity of differential binding affinities of the modulator to various reaction intermediates, are components of the theory of mechanism-based activation introduced below. This section presents statistical physics principles for the quantification and characterization of the effects of mechanism-based enzyme activators on protein local degrees of freedom and free energy differences in the catalytic mechanism, which will subsequently be related to experimentally measurable effects on enzymatic catalytic efficiency.

Computational studies of allosteric modulation^26,27^ employ methods such as dynamics simulations to sample protein global degrees of freedom, including domain motions. These global degrees of freedom are conveniently represented in terms of principal components. The principal components of the protein are the dominant (leading) eigenvectors of the covariance matrix (or normal modes) of the fluctuations in phase space degrees of freedom – namely the set of phase space positions and moments (*r*_*ix*_, *r*_*iy*_, *r*_*iz*_; *p*_*ix*_, *p*_*iy*_, *p*_*iz*_) of all the atoms *i*, which for simplicity can be expressed in cartesian coordinates. While the principal components (PCs) on both position and momentum phase space variables can be considered, the principal componentss on position variables are the relevant ones here. These are linear combinations of individual atomic degrees of freedom. We have the following expressions for principal componentss obtained from simulations (e.g., Monte Carlo or molecular dynamics) of global protein motions:

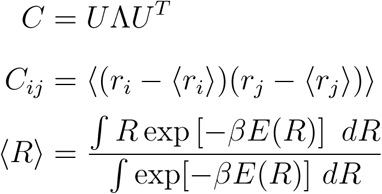

where *U* denotes the matrix of eigenvectors.

The principal coordinates can be expressed in other coordinate systems, in particular the set of all bond lengths, bond angle and dihedral angles under the *Z*-matrix representation, through a simple coordinate transformation – or they may be calculated. Hence, the principal componentss for a given protein can be related to (a large number of) local degrees of freedom. The fluctuations in these degrees of freedom obtained from (sufficiently long) molecular dynamics or Monte Carlo simulations include contributions from the conformational states involved in allosteric transitions.

By contrast, the degrees of freedom affected by mechanism-based activators are local, not principal components. They correspond to backbone (and associated side chain) degrees of freedom for specified subsets of residues, such as flexible loops.

### A. Modulation of local degrees of freedom

By inducing changes in the probability distributions of local active site degrees of freedom, the differential binding affinity of A to the different intermediates in the catalytic mechanism induce effects on the free energy differences in that mechanism which were considered in the models above. Here we consider how the ΔΔ*G*’s may be induced structurally in terms of active site conformational degrees of freedom. Below, we will consider the effect of arbitrary concentrations of a modulator *A* on activity given specified *K*_*di,A*_’s of modulator binding to different intermediates,

The impact of a modulator A on the free energy differences relevant to an enzyme’s catalytic mechanism can be represented in terms of a potential of mean force that alters the probabilities of various receptor states in the conformational ensemble, and hence alters the free energy differences. We write the DOF of the protein receptor as follows:

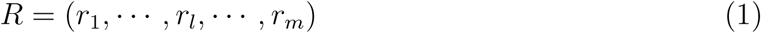

where the coordinates (*r*_1_, …, *r*_*l*_) highlighted are those whose conformational distributions change significantly in the presence of modulator A.

The potential of mean force induced by A on the receptor-ligand complex (RL) can be written

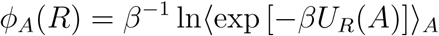

where *U*_*R*_ denotes the interaction potential between the receptor and modulator. As in the case of allosteric ensemble theory, it is possible to write the above expression in terms of dominant states in the conformational distributions of the local degrees of freedom, but for simplicity we do not do so explicitly here.

Because we are interested in the effects of displacements of the specified local degrees of freedom from nominal, favored conformations, we write the potential of mean force in terms of deviations from such a nominal conformation *R*_0_ as follows:

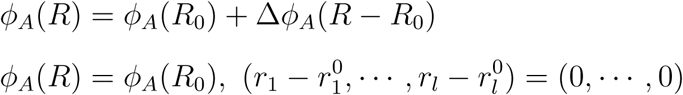

where the latter equation indicates that the potential of mean force only changes due to deviations of the specified local DOF, not due to other receptor motions like the global conformational changes relevant to allosteric activation.

Let 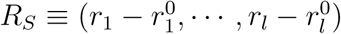. Assume a harmonic form of the change in the potential of mean force 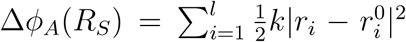. Note that 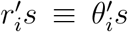 can denote dihedrals of backbone DOF instead of cartesian coordinates; in this case, we can set the units of *k* properly and restrict 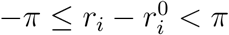.

Typically, mechanism-based activators may alter the distributions of local degrees of freedom whose conformations change during the course of the reaction, which leads to tradeoffs in the ΔΔ*G*′*s* for various reactions steps upon stabilization of one such conformation (see below). This is a critical distinction from allosteric activation, where there is no conformational change specific to certain steps in the reaction.

For example, activators of sirtuin enzymes that do not possess an allosteric site^18,19^ have been shown crystallographically to induce changes in the conformation of the so-called flexible cofactor binding loop in sirtuin enzymes. This loop changes conformation after the first chemical step of the reaction, namely the cleavage of nicotinamide from the NAD+ cofactor. In these cases, the local degrees of freedom above are the backbone degrees of freedom in the flexible cofactor binding loop.

### B. Induction of ΔΔ*G*’s in catalytic mechanisms

The free energies of L binding in the presence and absence of A can then be written in terms of exponential means of *ϕ*_*A*_(*R, L*) and *ϕ*_*A*_(*R*), providing an expression for the modulator’s effect on the free energy difference solely in terms of this potential of mean force. In the case of substrate binding free energy, for example, we have in the absence of *A*:

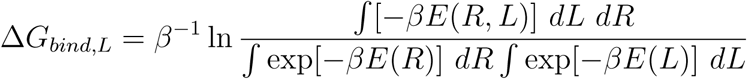

whereas in the presence of *A*:

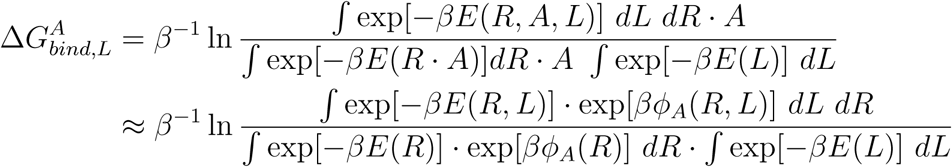

Each term above is the Δ*G* of interaction between the respective complex and *A*.

Then it can be shown that

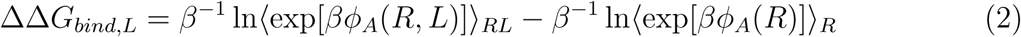

because

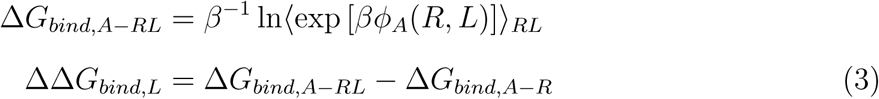

and

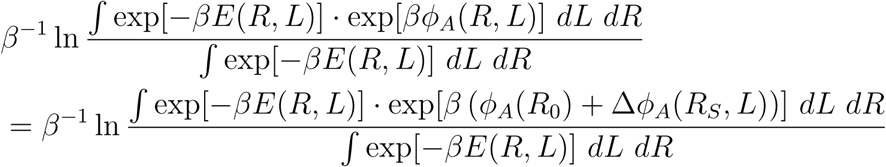

where *R, L* denote the receptor and ligand, respectively.

These expressions can be extended to free energies of activation. For such free energy differences of activation or chemical step equilibrium, instead of R, L in the expressions above, we use R with the subset of *r*_*i*_s that parameterize reaction coordinate constrained to specific values in the reactant and transition state geometries, in the numerator and denominator of 4 above. The free energy difference is hence expressed in terms of one partition function in numerator and denominator but with specified constraints on the atomic coordinates (reactant and transition state geometries) which change along the reaction coordinate.

In the models below, the *K*_*d,A*_’s of a modulator *A* to the various reaction intermediates induce these ΔΔ*G*’s. We note that allosteric modulation theory is generally not concerned with the full set of such free energy differences since the modulator typically, and design of modulators that have prescribed effects on the local degrees of freedom that induce these free energy changes is generally not possible.

## III. MECHANISM-BASED ACTIVATION OF TWO SUBSTRATE, TWO PRODUCT ENZYMATIC REACTIONS

A prerequisite for enzyme activation is that the modulator must co-bind with substrates. Within the context of enzyme inhibition, two modes of action display this property: noncompetitive and uncompetitive inhibition. Noncompetitive inhibitors bind with similar affinities to the apoenzyme and enzyme-substrate, enzyme-intermediate or enzyme-product complexes whereas uncompetitive inhibitors bind with significantly lower affinity to the apoenzyme. Both are specific examples of the more general notion of a mixed noncompetitive modulator that co-binds with substrates. Mixed mode modulation of enzymes by small molecules is typically not modeled in a steady state framework. However, this is essential for the study of mechanism-based activation because only steady-state models provide an accurate representation of activity.

We now explore how activation may be modeled by augmenting steady-state kinetic models for certain classes of enzymatic reactions, including those regulating flux through metabolic pathways, to include putative mechanism-based activators (A) that can bind simultaneously with substrate and product. Because such classes of activators have been reported for sirtuin enzymes, we carry out a representative analysis for enzymatic reactions that have two substrates and two products. This is also one of the simplest possible extensions of standard one substrate, one product reversible enzymatic reactions. Such reactions have the following steady-state rate law at saturating [*S*_2_] (denoting *S*_1_ by *S* and *P*_1_ by *P* for simplicity):

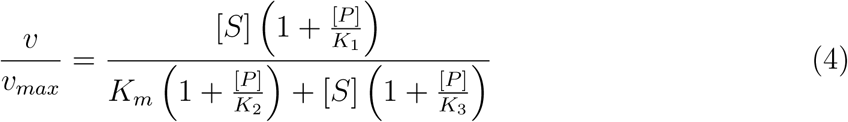

The basic steady-state kinetic model for such reactions was presented in^21,28^. Since the augmented kinetic model in the presence of a mechanism-based modulator depends on the expressions for each of the intermediate species in the enzymatic reaction in terms of the rate constants and the concentrations of substrate and product, the analysis below is presented in terms of the coefficients *c*_*ij*_ in these expressions, where *i* denotes the index of the reaction intermediate and *j* denotes the term in the polynomial expansion of the concentration of this reaction intermediate on substrate/product concentrations and products thereof. These coefficients are presented in the Appendix for the example of sirtuins.

Fig. 2 depicts the reaction diagram for mechanism-based activation. Note that only the top and front faces of this cube are relevant to the mechanism of action of the previously proposed competitive inhibitors of base exchange and deacylation. At any [A] there exist apparent values of each of the rate constants in the reaction mechanism. These are denoted by app in the Figure. There are also corresponding app values of the steady state, Michaelis, and dissociation constants.

**FIG. 1.**
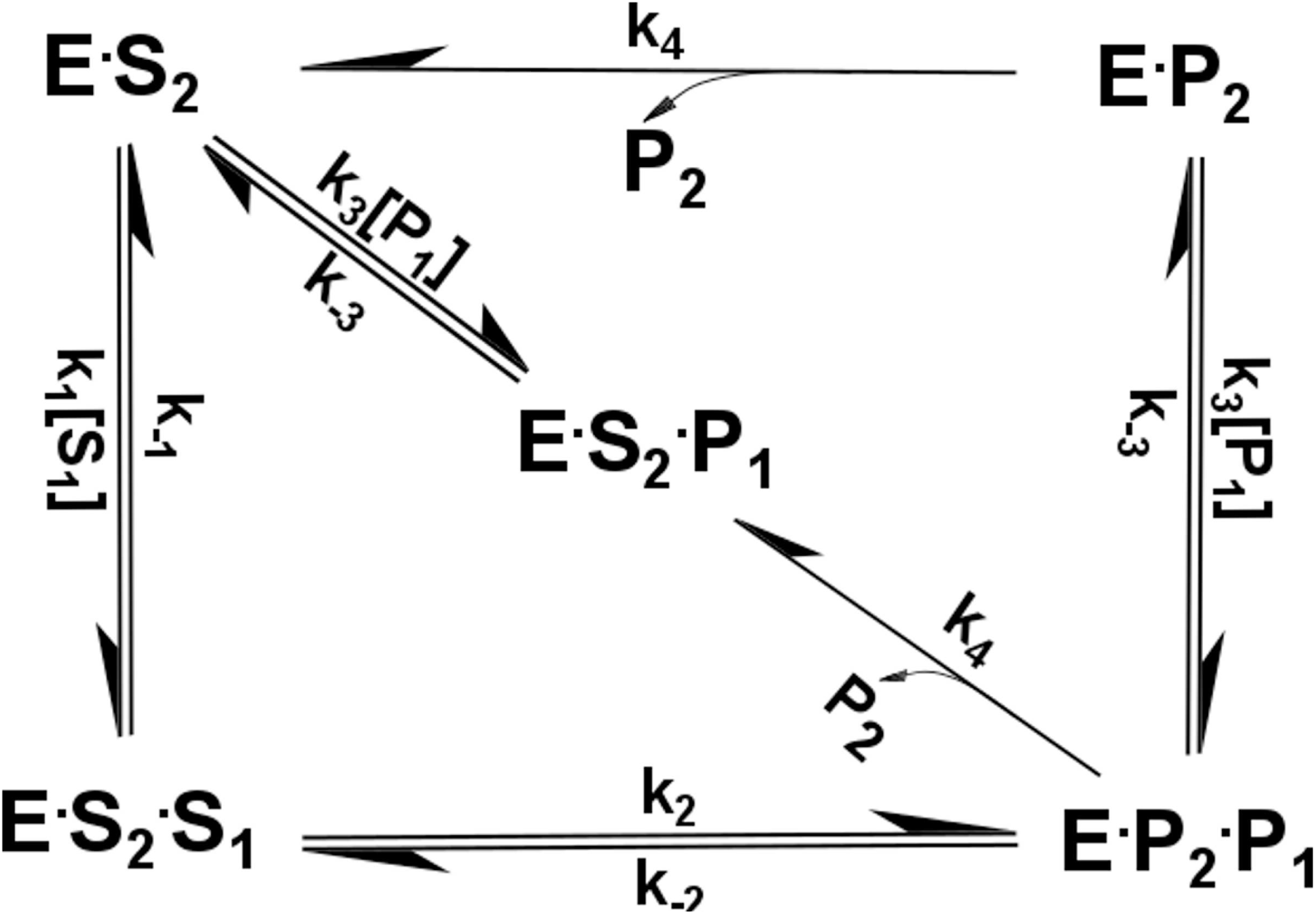
Basic reaction scheme for two substrate, two product reactions where one substrate is present at saturating concentration.

**FIG. 2.**
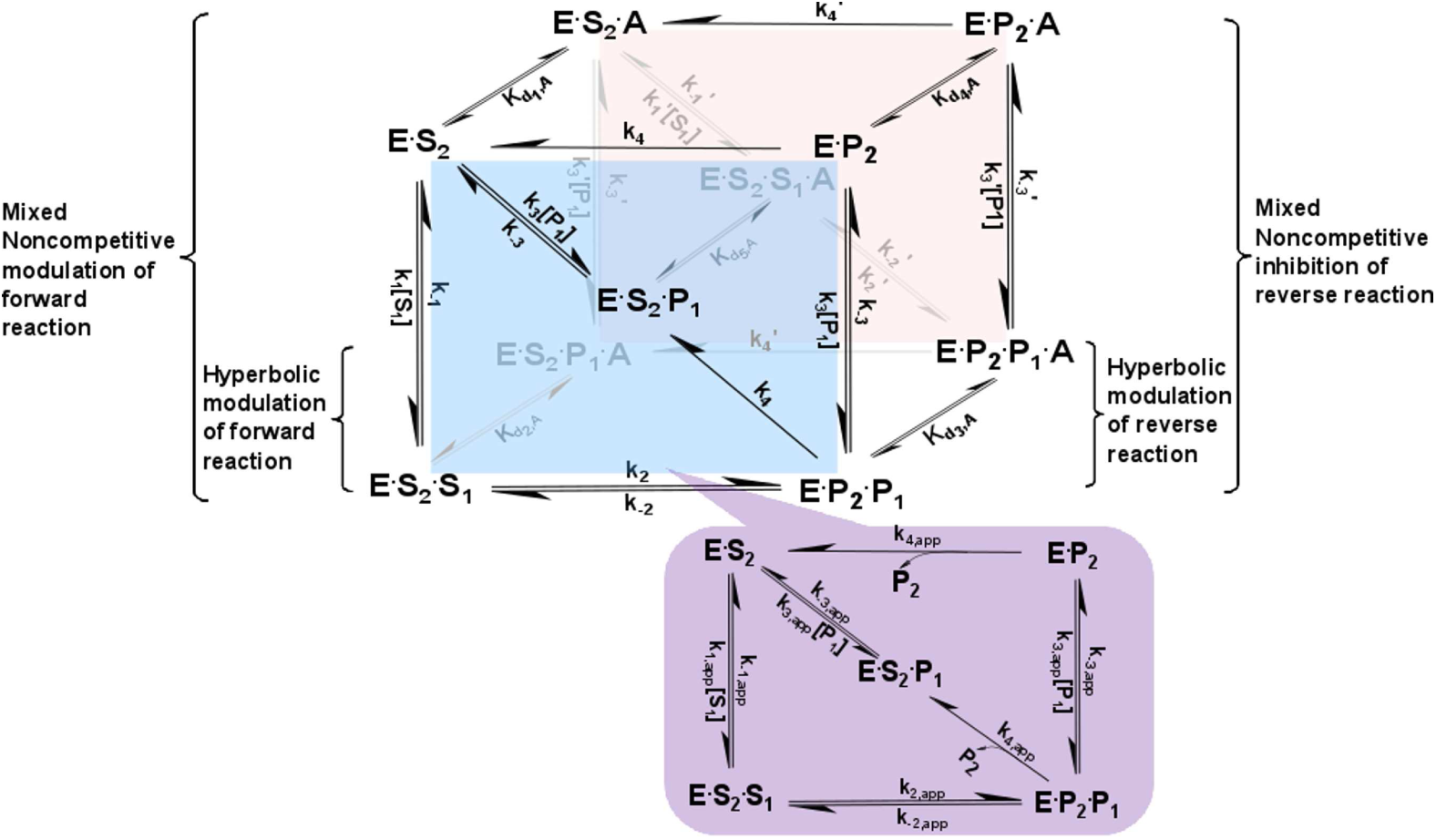
Reaction network for mechanism-based enzyme activation of a two substrate, two product enzymatic reaction. The front face of the cube (blue) depicts the salient steps of the sirtuin reaction network in the absence of bound modulator. The back face of the cube (red) depicts the reaction network in the presence of bound modulator (denoted by A). Each rate constant depicted on the front face has an associated modulated value on the back face, designated with a prime, that is a consequence of modulator binding. In the absence of modulator, the reaction proceeds solely on the front face, whereas in the presence of saturating concentration of modulator, the reaction proceeds solely on the back face. The purple projected face is the apparent reaction network in the presence of a nonsaturating concentration of modulator. On this face, each rate constant is replaced by an apparent value, denoted by app. Mixed noncompetitive modulation of the forward and reverse reactions involves binding of A to the reaction species on the left and right sides of the front face, respectively, whereas hyperbolic modulation by A involves alteration of the rate constants for forward and reverse reactions (*k*_2_ and *k*_−2_, respectively). For small [A], the effect of the side and back faces of the cube on the apparent rate constants is modeled under a rapid equilibrium approximation, whereas a full steady state analysis is applied to the front face.

Since the magnitudes of the *K*_*d,A*_s or binding affinities (Δ*G*_*bind,A*_’s) of A do not directly affect the shape of the dose response curves and the maximum level of activation, the ratios of *K*_*d,A*_s and hence the relative binding affinities of the front and back face complexes (Δ*G*_*bind,A*_s) are the thermodynamic quantities of interest. In order to obtain predictions for the effect on 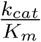 of a modulator with specified relative binding affinities for the complexes in the reaction mechanism – which is crucial to the mechanism of action of a potential activator – it is important to develop a model that is capable of predicting, under suitable approximations, the effect of a modulator with specified binding affinities on the apparent steady state parameters of the enzyme. For characterization of a known activator, one can carry out complete steady state system identification at saturating [A]^29^ to estimate the actual 7 back face rate constants in the presence of bound A (the rate constants designated by primes in the Figure).

The following notational mappings are used for convenience in the derivations: The mapping of the rate constants is as follows: 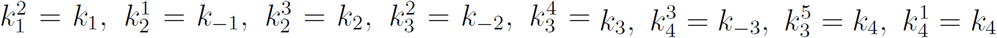. The mapping of the species notations is apparent from the figure.

Fig. 3 provides a basis for the derivation of the modulated expressions for the steady state constants of the enzymatic reaction. Note that the number of edges is n-1=9. Any 0th order term in [A] (which all contain a front face that is feasible alone) must contain five arrows from back to front face, just as any 5th order term in [A] must contain five arrows from front to back face. The *c*_*ij*_ in the Appendix are all multiplied by a common factor to provide the 0th order terms.

**FIG. 3.**
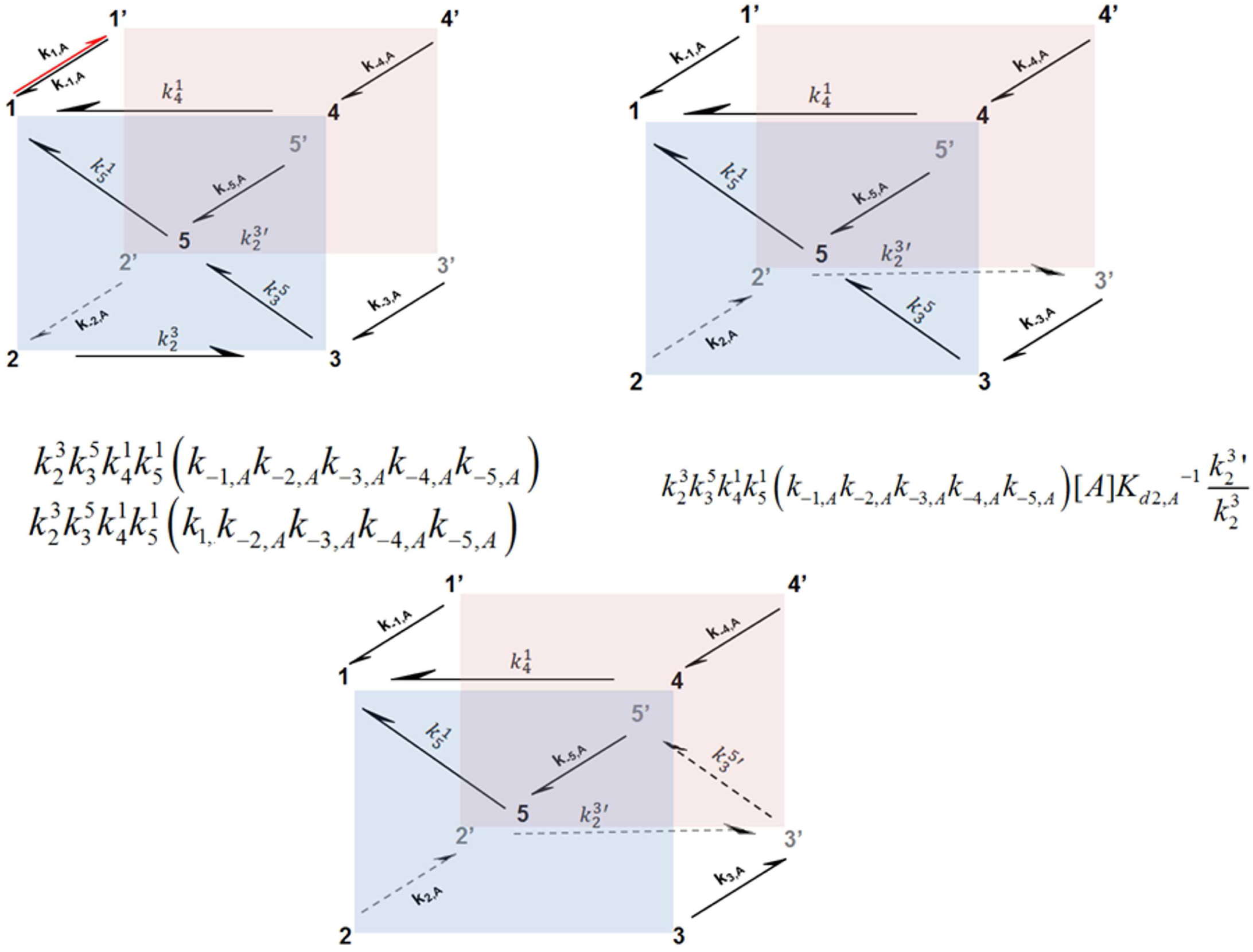
Graphical depiction of representative King-Altman terms [Representative King-Altman diagrams] in mechanism-based enzyme activation. Order refers to that in the concentration of activator, [A]. B. 0th-order diagram; C. Representative 1st-order term; D. Representative 2nd-order term.

Since the full steady state expression relating the original to the apparent rate constants has many terms containing products of additional side and back face rate constants, we use a simpler augmented kinetic model to in order to explore the prospects for mechanism-based activation of enzymes. Due to the scaling of the number of terms in the expressions, we apply the following approximation: *k*_*i,on*_, *k*_*i,off*_ ≫ *k*_*j*_, *i* ≠ 1, …, 5; *j* ≠ *i, on, j* ≠ *i, off*. Then, every nonnegligible term/diagram that is 1st to 4th order in [A] and that ends in a species on the front face can be constructed by an appropriate transformation of one of the above constant terms/diagrams involving moving an arrow from front to back face and flipping direction of side face arrows.

The purpose of the approximation is to consider the effect of A on kinetic parameters in terms of equilibrium biophysical properties of its binding to reaction intermediates, without needing to know the values of all rate constants. We arrive at simple definitions of the apparent Michaelis constant and other steady state constants for the reaction in terms of the original expressions for these constants and the dissociation constants for binding of A to the various complexes in the enzymatic reaction mechanism. This provides a minimal model with the least number of additional parameters required to model enzyme activation mechanisms. The rapid equilibrium segments model is introduced in order to consider the plausibility and biophysical requirements of mechanism-based activation based only on the free energy changes of the various species in the sirtuin reaction mechanism upon binding A. This model assumes the changes in species concentrations in the presence of A are determined by the *K*_*d,A*_’s in Fig. 2.

We illustrate the principles of the analysis with a specific derivation of the modulated expression for *K*_*m*_; analogous principles apply to other steady state constants. Details of the derivation and definitions of notations are provided in the Appendix. Following this derivation, we have

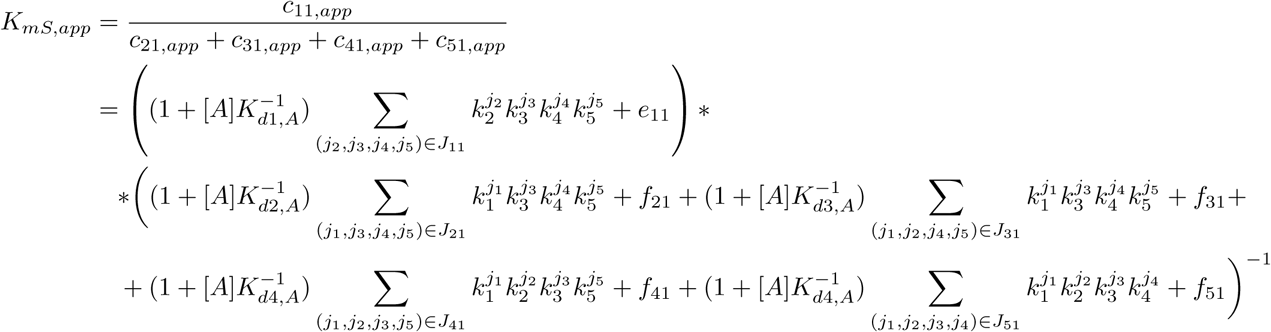

with

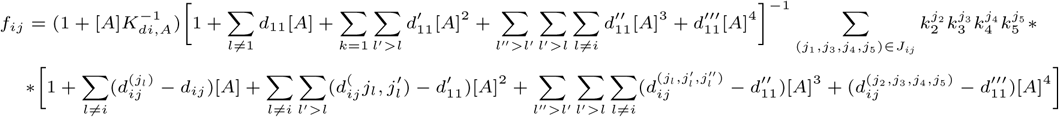

### A. Regimes of interest

Consider two regimes of interest:

1. *k*_*i,off*_ ≫ *k*_*i,on*_[*A*], *i* = 1, …, 5, where ≫ refers to a specified low/leading order approximation in [*A*] being valid (with error less than a specified threshold). Apply the above representation to arrive at the relevant approximation. Here, if we also have *ki, off* ≫ *kj, j* ≠ *off* (i.e., a weak binder), we can apply the “rapid equilibrium segments” approximation, which is discussed further below with reference to sirtuin enzymes. We assume it is true for purposes of analysis. It may especially be true for early hits in drug discovery. If it is not true, see regime 2 below.
2. *k*_*i,off*_ ≪ *k*_*i,on*_[*A*], *i* = 1, …, 5, where ≫ refers to a specified low/leading order approximation in [*A*]^−1^ being valid (with error less than a specified threshold). This case is discussed in the Supporting Information. We can always make the “rapid equilibrium segments” approximation in terms of the 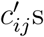 valid at sufficiently high [*A*] (choose [*A*] such that *k*_*i,on*_[*A*] ≫ *kj, j* ≠ *on*).

Note that the magnitude of [*A*]*/K*_*di,A*_ = *k*_*i,on*_[*A*]*/k*_*i,off*_ determines whether we are operating in the front face or back face regime. We are at symmetrical parts of the regimes when [*A*]*/K*_*di,A*_ (regime 2) = *K*_*di,A*_*/*[*A*] (regime 1). Whether we can neglect terms of certain orders in [*A*]*/K*_*di,A*_ depends on the magnitude of this ratio.

The equations above provide a model for dose-response properties of mechanism-based modulators at arbitrary [*A*]. This is discussed further below.

Similar principles can be applied to one substrate, one product enzymatic reactions. The Supporting Information displays the corresponding schematic for modulation of such reactions.

### B. Prediction of approximate dose-response curves for mechanism-based activators

The approximate expressions derived in the Appendix for the dose-response properties of mechanism-based activators, can be applied if the front and back face rate constants (*k*_*i*_ and 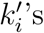) are all known. These expressions are fifth-order in [*A*] for the two-step enzymatic reactions considered and third-order in [*A*] for the one-step reactions. The complete set of unmodulated and modulated rate constants for a given modulator can be obtained through experimental system identification methods in the presence of absence of modulator, through computational methods given crystal structures of the cocomplexes of modulator with the various reaction intermediates (of the type solved, e.g., for sirtuins in reference^30^), or a combination of both. As such, the theory presented herein can provide approximations to the dose-response behavior of mechanism-based enzyme activating compounds, which are fundamentally different from those of traditional inhibitors or allosteric activators.

## IV. EXAMPLE: MECHANISM-BASED ACTIVATION OF SIRTUIN ENZYMES

To render the analysis concrete, we now consider application of the formulation above to the specific example of sirtuin enzymes and apply simplifying approximations. These enzymes have an acylated protein substrate (Ac-Pr) and cofactor NAD+. Mammalian sirtuins (SIRT1-SIRT7) are a family of nicotinamide adenine dinucleotide (NAD+)-dependent protein deacylases and play critical roles in lifespan and aging related disease^31^. Sirtuins require the cofactor NAD+ to cleave acyl groups from lysine side chains of their substrate proteins, and producing nicotinamide (NAM) as a by-product. The mechanism of sirtuin-catalyzed deacylation is depicted in Fig. 4. Of the seven sirtuin enzymes, only one (SIRT1) possesses a known allosteric site. Activation by small molecules of some of the other six sirtuins, which do not have allosteric sites, is the subject of intense basic and translational research interest, but no framework for the design of such activators exists. A preferred general strategy for activation of sirtuins, as depicted in Fig. 4, would be to lower the *K*_*m*_ for the NAD+ cofactor. Unlike allosteric activation that increases affinity for the protein substrate of the enzyme, this approach could be applicable to multiple sirtuins and substrates.

**FIG. 4.**
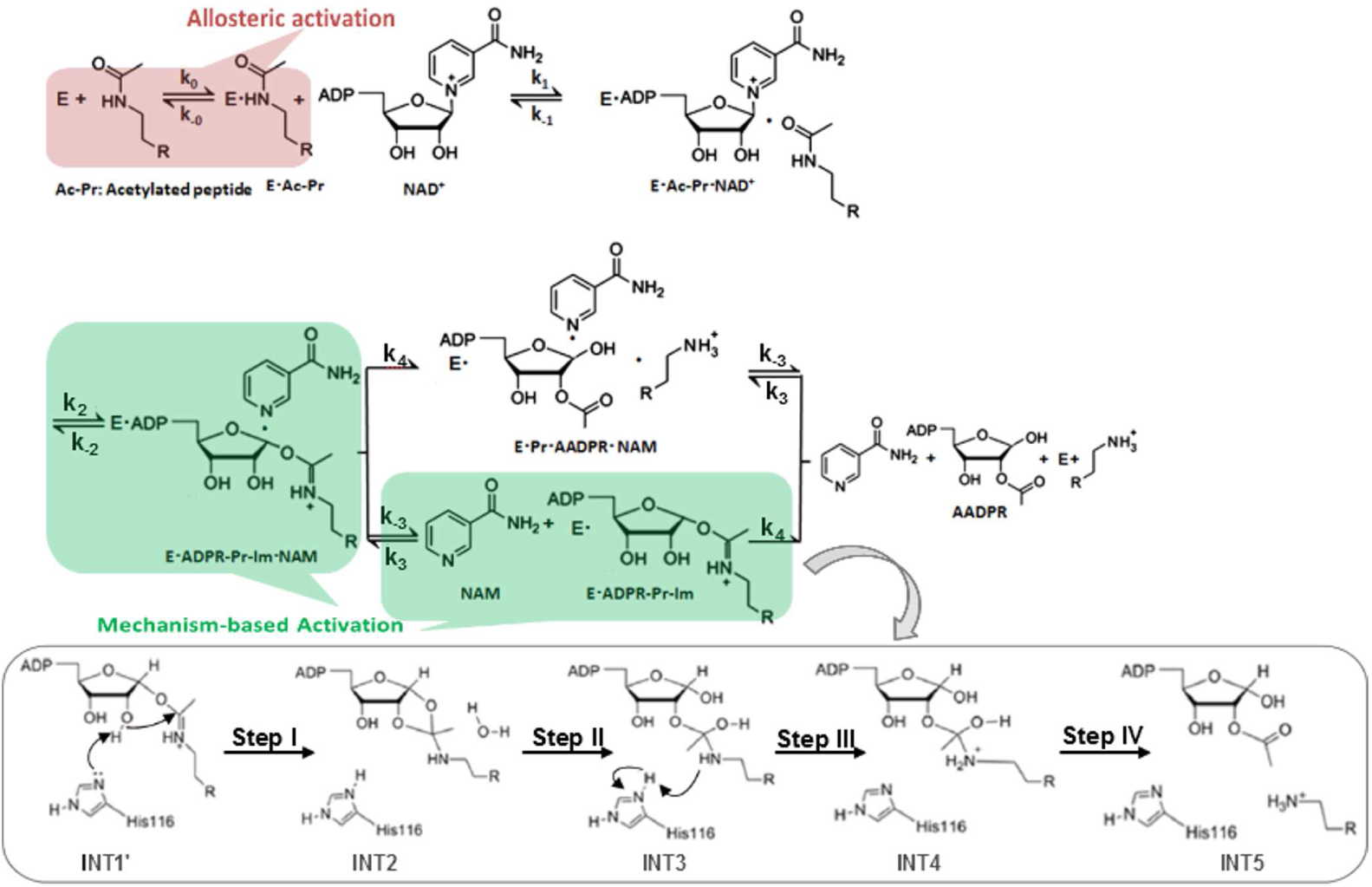
Chemical mechanism of sirtuin-catalyzed deacylation and modes of sirtuin activation. Following sequential binding of acylated peptide substrate and NAD+ cofactor, the reaction proceeds in two consecutive stages: i) cleavage of the nicotinamide moiety of NAD+ (ADP-ribosyl transfer) through the nucleophilic attack of the acetyl-Lys side chain of the protein substrate to form a positively charged O-alkylimidate intermediate (O-AADPR), and ii) subsequent formation of deacylated peptide. For simplicity, all steps of stage ii as well as AADPR + Pr dissociation are depicted to occur together with rate limiting constant *k*_4_. The schematic highlights mechanism-based activation through NAD+ *K*_*m*_ reduction rather than the peptide *K*_*d*_ reduction that known allosteric sirtuin activators elicit.

As mentioned before, several compounds have been reported in the literature as being activators of sirtuins other than SIRT1^18,20^. In addition, sirtuins whose preferred substrates are long chain acylated proteins were reported to have their activities on shorter chain acyl groups increased by certain fatty acids, which further corroborates the feasibility of nonallosteric sirtuin activation^19^. The theory presented in this paper provides a unifying framework under which such compounds can be characterized and hits can be evolved into leads.

Overexpression or activation of multiple mammalian sirtuins – including SIRT1,SIRT2, SIRT3, SIRT6 and SIRT7 – has been demonstrated experimentally to extend lifespan, alleviate age-related diseases and/or reverse the symptoms of age-related health decline^3,13,14,20,32^. The following small molecules have been reported to activate mammalian sirtuin enzymes that are involved in the regulation of healthspan and lifespan, and that lack allosteric sites:

- long-chain fatty acids: SIRT6 (deacetylation)^19^
- multiple quinolone derivatives: SIRT5 (desuccinylation), SIRT6 (deacetylation)^18^
- honokiol: SIRT3 (deacetylation)^20^

In the case of fatty acids and quinolone compounds, structural studies have demonstrated the binding occurs at the active site and induces conformational changes that improve catalytic efficiency. The catalytic efficiency was increased over 30-fold for fatty acids and several fold for quinolone compounds. Structural studies of the binding of resveratrol, in the same chemical family as honokiol, suggest that honokiol binding also occurs at the active site. Structural studies of quinolone binding demonstrate the local conformational changes (discussed in Section 2) in the active are induced. Moreover, *K*_*m,NAD*+_ can vary significantly for substrates with different acyl chain lengths^31^, demonstrating the potential to modulate catalytic efficiency based on local active site conformational modulation.

Here, we have the following mappings of species at saturating peptide and unsaturating cofactor: 1 : *E*.*Ac* − *Pr*, 2 : *E*.*Ac* − *Pr*.*NAD*; 3 : *E*.*ADPR* − *Pr* − *Im*.*NAM* ; 4 : *E*.*ADPR* − *Pr* − *Im*; 5 : *E*.*Ac* − *Pr*.*NAM*, where ADPR-Pr-Im is the alkylimidate reaction intermediate. *S*_1_ is hence NAD+, *S*_2_ is Ac-Pr, *P*_1_ is NAM, and *P*_2_ is DeAc-Pr. Unsaturating cofactor is the condition of greatest medicinal interest because depletion of NAD+ cofactor is a leading cause of age-related health decline. In what follows, 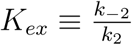.

Along with SIRT1, SIRT3 is one of the most important sirtuin enzymes involved in the regulation of healthspan^14^; hence we chose it as a subject of study. Honokiol (HKL) has been reported as a SIRT3 activator^20^. Because SIRT3 does not share the allosteric activation site of SIRT1, we studied HKL as a potential hit compound for mechanism-based activation of SIRT3. Extensive additional studies by our group on the mechanism whereby HKL activates SIRT3 are reported in reference^33^.

### A. Methods: modulation of SIRT3 activity by honokiol

#### Docking of HKL to the SIRT3.Ac-Pr.NAD+ substrate complex by Auto-Dock Vina

Molecular dockings on QM/MM optimized structure of SIRT3 have been carried out by AutoDock–Vina. AutoDock–Vina repeatedly docks each ligand to the target applying several search algorithms like global search implementing simulated annealing genetic algorithm, local search algorithm hybrid global–local search algorithm and also Lamarckian genetic algorithm and iterated local search, i.e. genetic algorithm with local gradient. AutoDockTools have been used to prepare the receptor PBQT format adding polar hydrogen and assigning Gasteiger charges to all its atoms as well as identifying the coordinates of the target box to enclose QM/MM optimized SIRT3.Ac-Pr.NAD+ substrate for global docking with a cubic grid box of 126×126×126 point dimensions of 0.5 Å resolution. Prior to docking, Honokiol structure was geometry optimized in gas phase by Hartree-Fock method with 6-311G+ basis.

#### Experimental methods

Please see Supporting Information for details of experimental methods.

### B. Docking of honokiol to SIRT3 and microscale thermophoresis

HKL was docked to the SIRT3.Ac-Pr.NAD+ substrate complex. As depicted in Fig. 5 and 6, HKL can cobind with substrates, at a site near the active site, as required for mechanism-based enzyme activators. HKL demonstrated high affinity (−5.8 kcal/mol) binding to a pocket next to the active site, and is predicted to engage in H-bonds with Glu177 of the flexible cofactor binding loop which will induce structural changes due to loop flexibility, consistent with the theory of mechanism-based activation above. These direct interactions between HKL and the flexible cofactor binding loop are responsible for generating the potential of mean force *ϕ*_*A*_(*R*) derived in Section 2A.

**FIG. 5.**
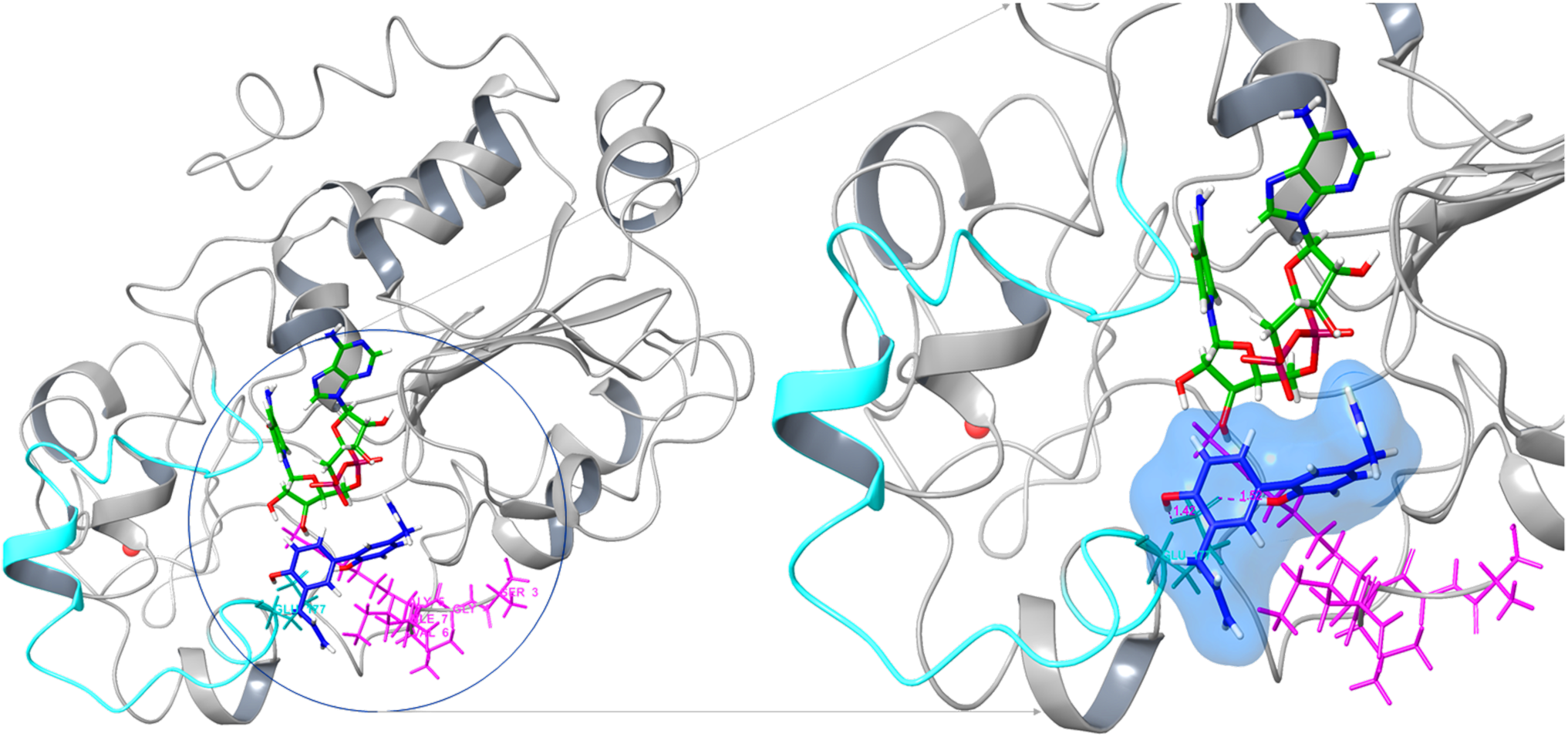
Docking of HKL to the SIRT3.Ac-Pr.NAD+ substrate complex: Binding mode of honokiol to the SIRT3 ternary reactants complex as predicted by docking. The predicted binding free energy is -5.8 kcal/mol. The SIRT3 protein is represented in its secondary structure colored in white. The acetylated ACS2 peptide is represented in stick model, and is depicted in pink. The Honokiol ligand is shown in blue sticks. The Honokiol molecule forms H-bonds with Glu 177 which is present in the cofactor binding loop (highlighted in cyan color in the secondary structure of SIRT3) of the SIRT3 protein.

**FIG. 6.**
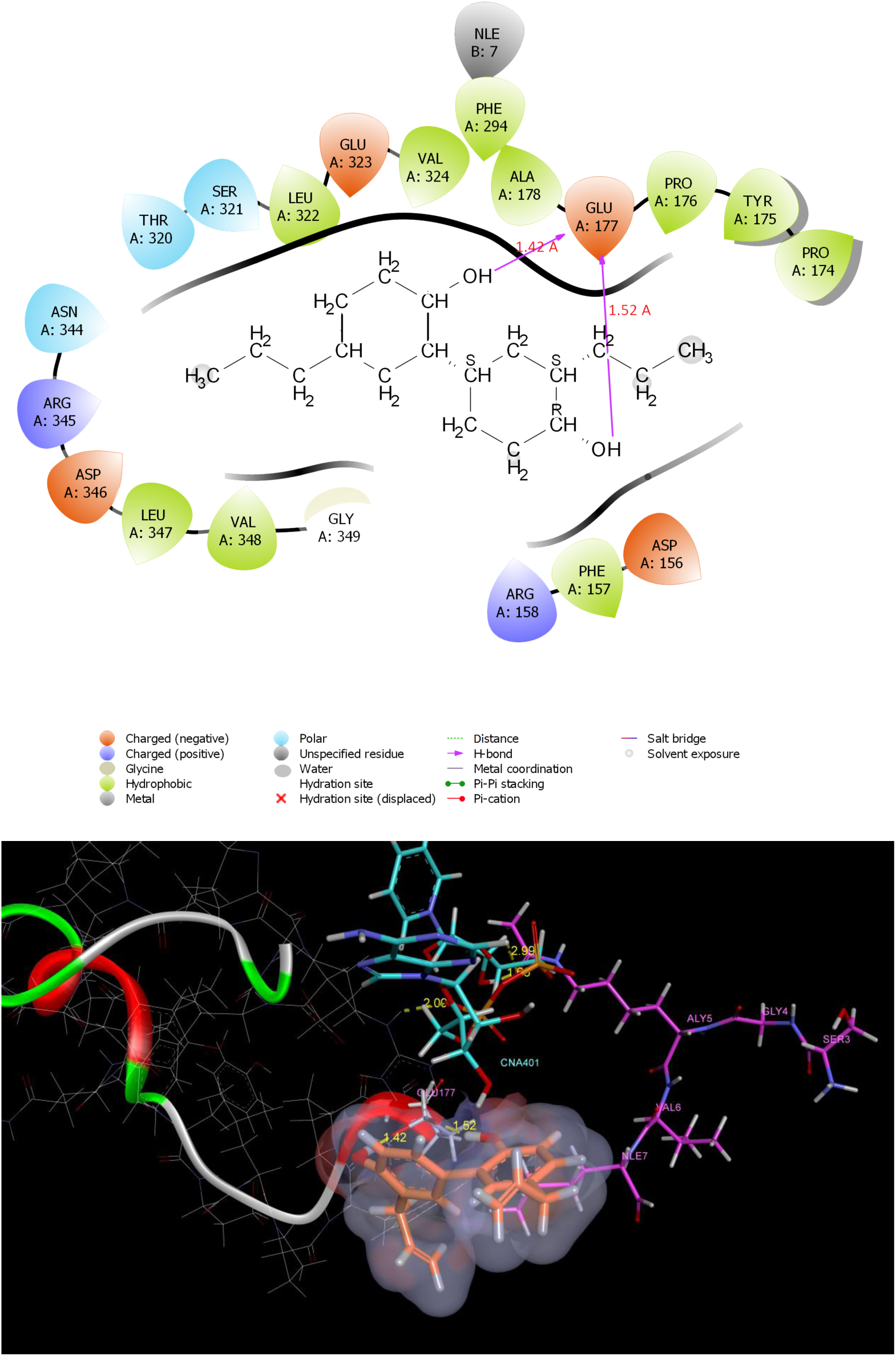
Docking of HKL to the SIRT3.Ac-Pr.NAD+ substrate complex: Ligand interaction diagrams of honokiol:SIRT3 ternary reactants complex binding as predicted by docking. **Top:** 2D ligand interaction diagram. **Bottom:** 3D ligand interaction diagram highlighting H-bonding. The Honokiol ligand binds with the SIRT3 protein at Glu 177 position and forms two hydrogen bonds with two of its hydroxyl groups present in it. The Glu 177 is present in the cofactor binding loop of SIRT3 protein.

Moreover, microscale thermophoresis measurements on carba-NAD+ indicate cofactor, substrate and honokiol all cobind with a slight decrease in NAD+ binding affinity. Reference^33^ reports the experimental characterization of SIRT3 modulation by honokiol for another substrate (MnSOD), in which case the binding affinity of NAD+ is increased by honokiol binding.

### C. Non-steady state activation of SIRT3 by honokiol

Fig. 7 depicts how HKL activates human SIRT3 under non-steady state conditions, including the dependence of activation on enzyme concentration and reaction time. Due to the sensitivity and convenience of the fluorescence-based assay for SIRT3 deacetylation activity on the fluorolabeled p53 substrate, this assay can be employed using the reaction conditions depicted in this Figure in the context of screening of libraries of drug-like compounds to identify hit compounds for mechanism-based activation of the SIRT3 enzyme.

**FIG. 7.**
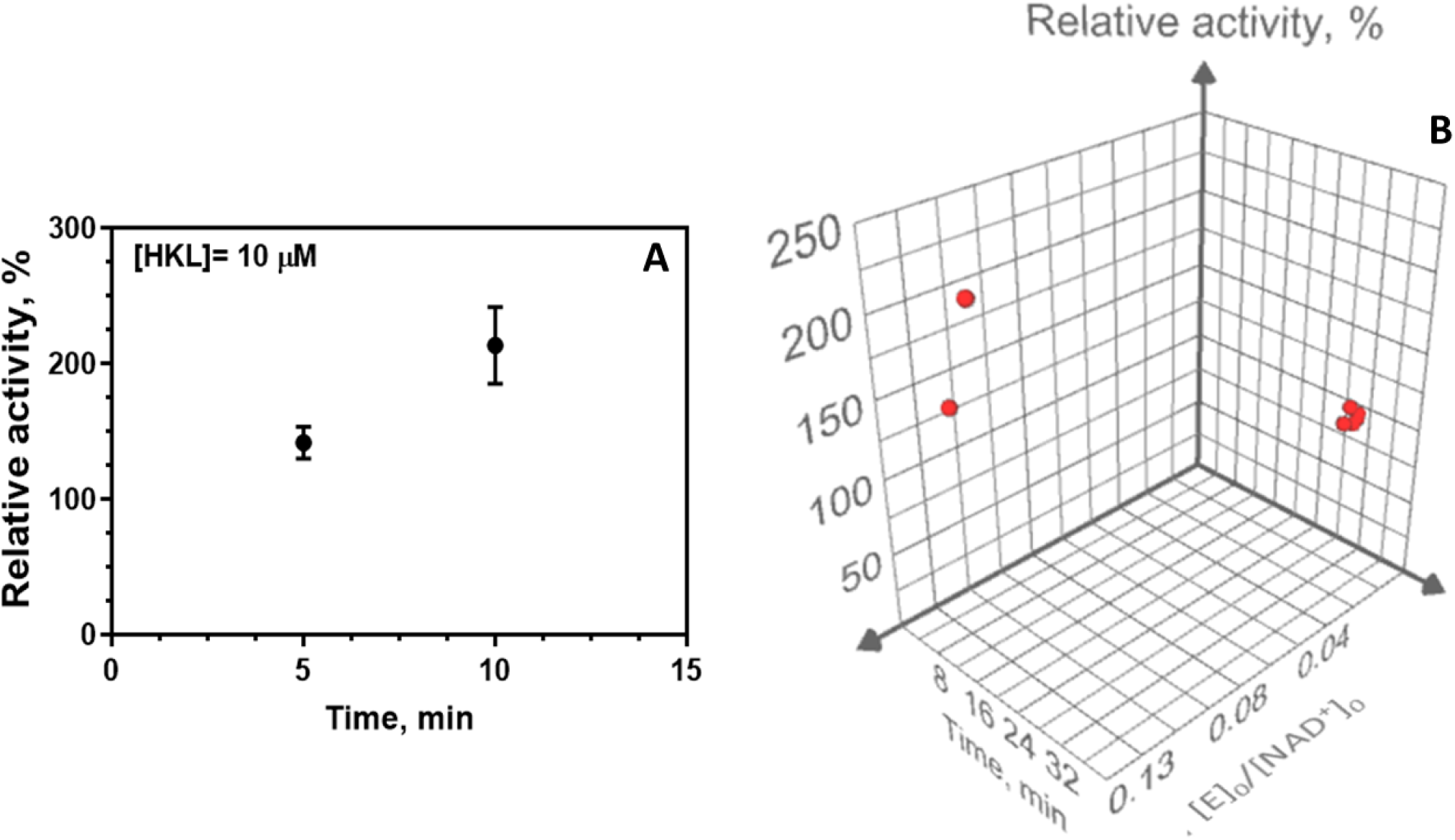
Non-steady state activation of human SIRT3 by honokiol. (A) Dependence of activation on time (fixed enzyme concentration and [*E*]_0_*/*[*NAD*+]_0_ = 0.122, using 10uM NAD+ and 10uM fluorolabeled acetylated p53 peptide substrate). (B) Dependence of activation on enzyme concentration and time. [HKL]=10uM.

In reference^33^, the effect of HKL on the rate of SIRT3-catalyzed deacetylation of the physiologically important MnSOD substrate was studied using a label-free assay. It was shown that the initial rate of SIRT3-catalyzed deacetylation of this physiologically important substrate increases in the presence of HKL, whereas the steady state rate does not increase. In the next section, we study steady state modulation of SIRT3-catalyzed deacetylation of the fluorolabeled p53 substrate.

### D. Steady state modulation of SIRT3 by honokiol

The results of steady state characterization of the effects of saturating [HKL] on SIRT3catalyzed deacetylation of the fluorolabeled p53 substrate are presented in Figs. 8,9. In these experiments, [*E*]_0_*/*[*NAD*+]_0_ *<<* 1 and reaction conditions were chosen to ensure only the steady state phase is measured.

**FIG. 8.**
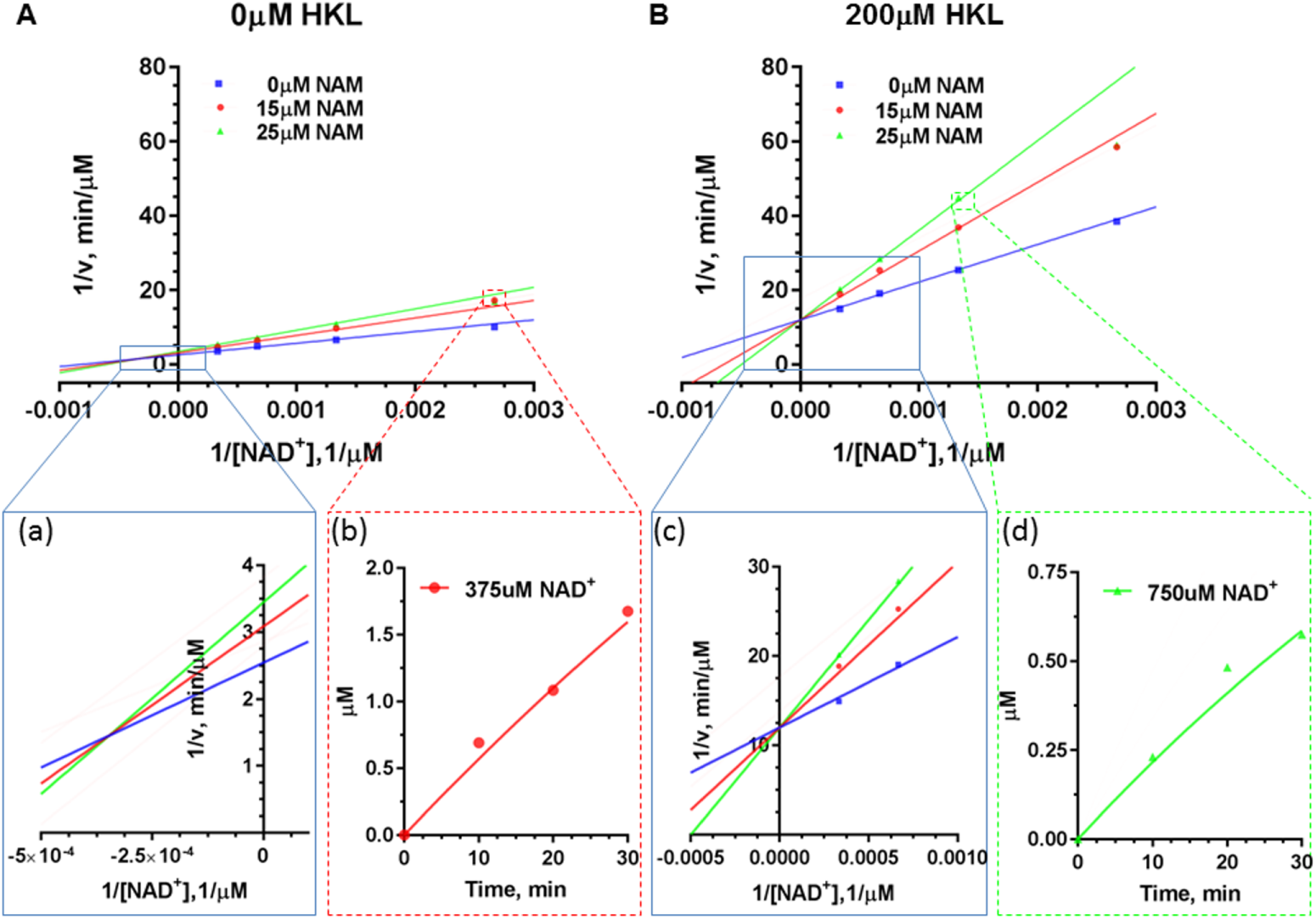
Double reciprocal plots for deacylation initial rate measurements of saturating substrate peptide (FdL2) in the presence of different concentrations of NAM with (A) 0 uM HKL; (B) 200 uM HKL. The enlargement of the intersection points were provided as insets (a) and (c). The time series plot of mM product formed vs. time are provided as insets (b) and (d).

**FIG. 9.**
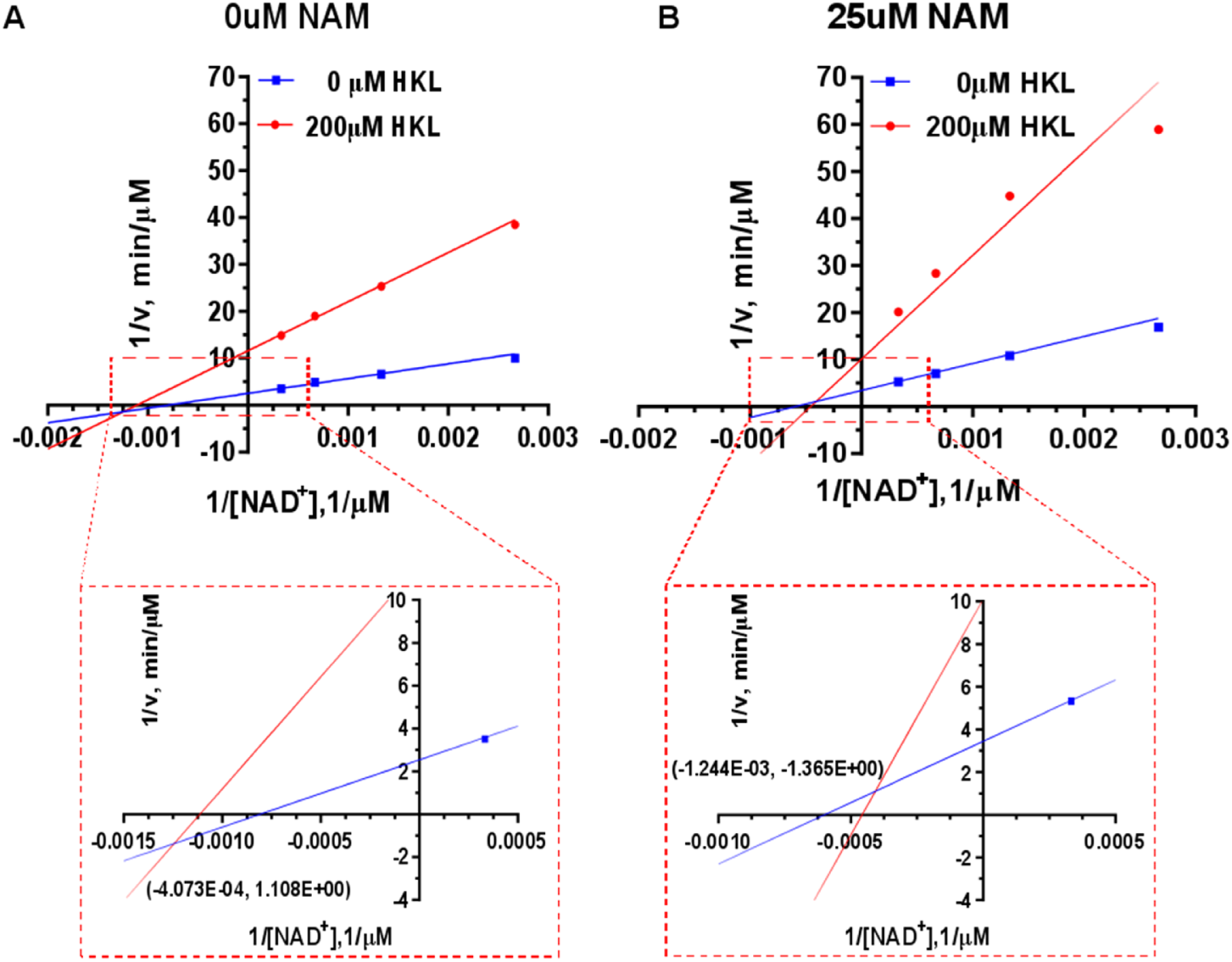
Double reciprocal plots for deacylation initial rate measurements of saturating substrate peptide (FdL2) in the absence and presence of Honokiol with (A) 0 uM NAM; (B) 25 uM NAM. The zoom in of intersection point with x, y values are provided as insets.

Expressions for apparent values of all steady state parameters introduced in equation (4) (i.e., modulated versions of constants (*v*_max_, *K*_*m,S*_, *K*_1_, *K*_2_, *K*_3_) in the presence of a given [A] can now be derived for this enzyme. In the following, several types of approximations will be invoked in addition to that applied above:

1. rapid equilibrium segments approximation: a)

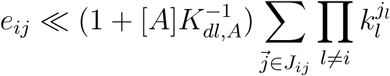

b)

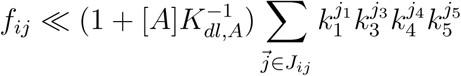
2. *k*_4_(1 + *K*_*dl,A*_) ≪ *k*_*j*_(1 + *K*_*dl*_’,*A*), *j*≠4, *l* = 1, …, 5
3. *k*_−3_(1+*K*_*dl,A*_) ≫ *k*_*j*_(1+*K*_*dl*_*1,A*), *j* ≠ 2, *l* = 1, …, 5 (rapid P dissociation; this typically holds for sirtuins)

In what follows, please note we use the original notations for the rate constants as shown in Figs. 1 and 4.

Under these additional approximations, at low [A] we assume that the changes in each of the rate constant products in *c*_*ij*_ and *c*_*i′j′*_, *i′* = *i* are the same and linear in [A]:

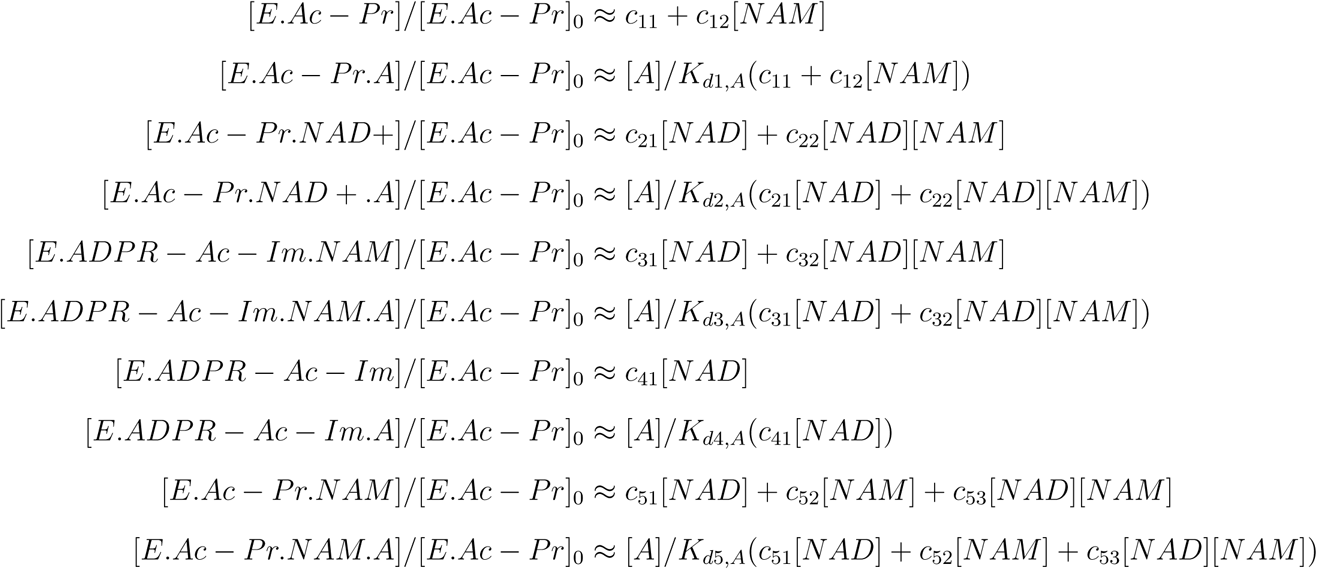

Under these additional approximations, at low [A], the rapid equilibrium segments expressions for the various steady-state species concentrations are then as follows (please note that in order to model the dose-response behavior of the modulator, the full equations presented in Section 3 and the Appendix are needed):

- Effect on 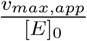:

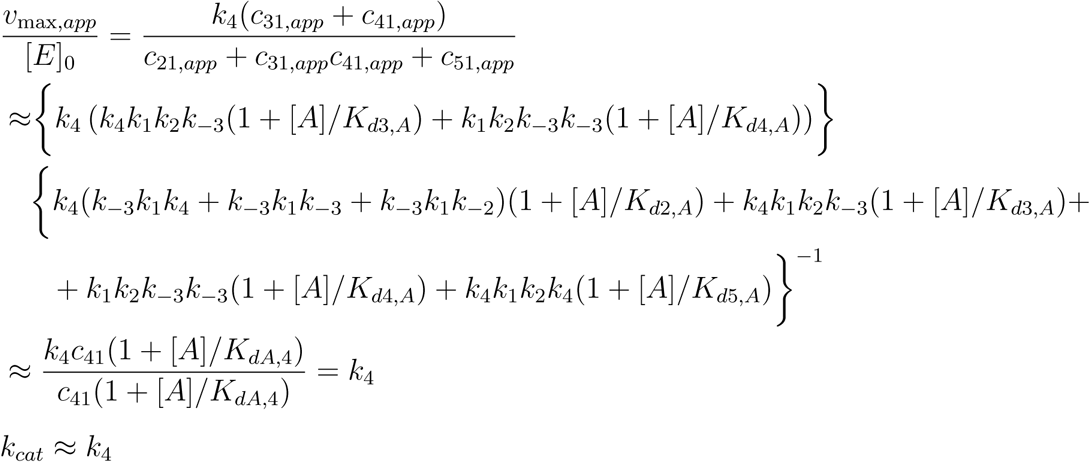
- Effect on *K*_*mNAD*_+,*app*:

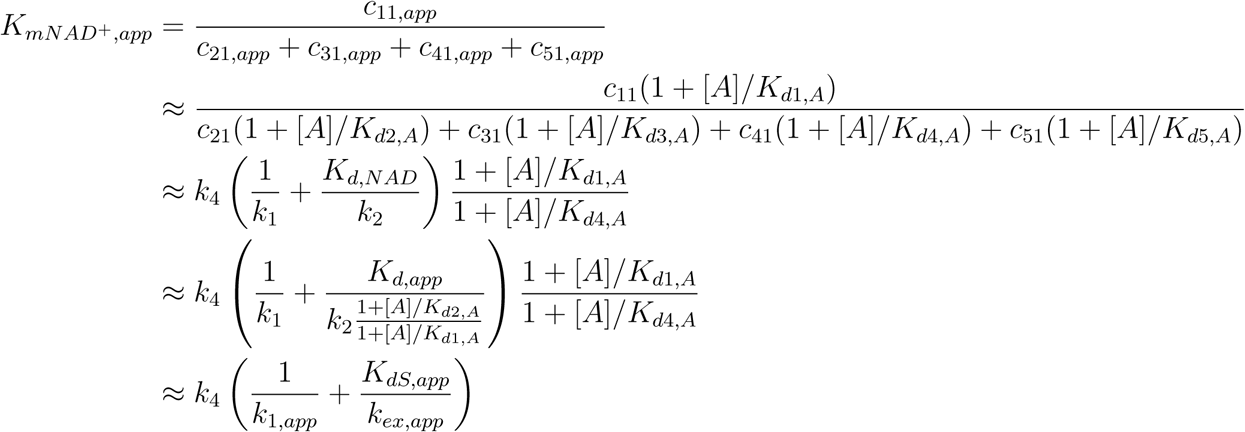
- Effect on *α*_*app*_ and *α*_*app*_*K*_*mNAD*_+,*app* Recall that *α* provides an estimate of the ratio of the dissociation and Michaelis constants for S.

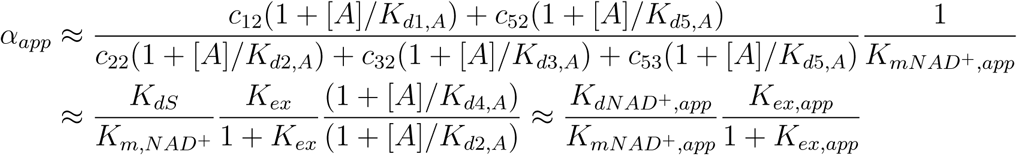
- Effect on *K*_3,*app*_:

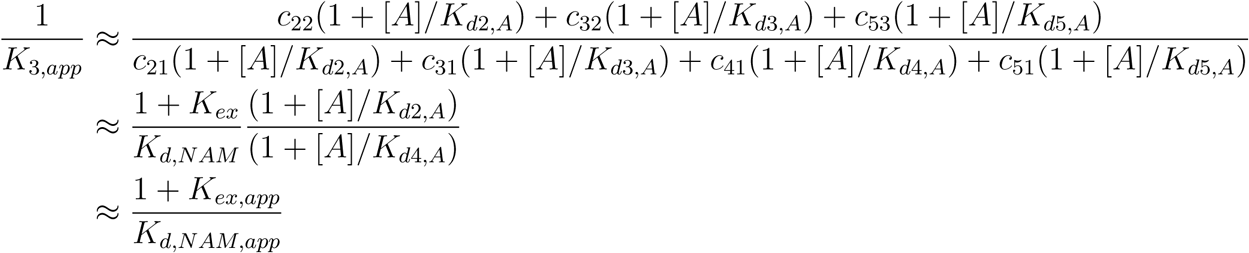
- Effect on *K*_2,*app*_:

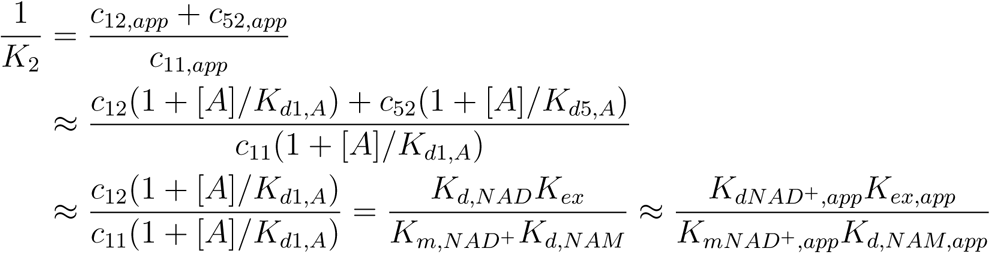
- Effect on *K*_1,*app*_:

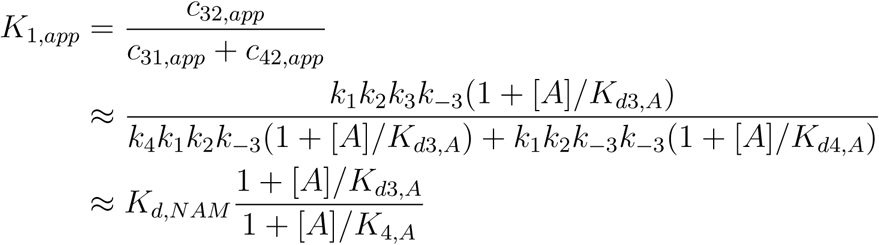

where 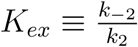.

Note that for competitive inhibition of the reverse reaction, unlike the noncompetitive modulation modes depicted in Fig. 2, interpretation of the apparent steady state constants in terms of the apparent dissociation and chemical equilibrium constants of the reaction mechanism are no longer valid since approximation (3) above does not hold for all *l*.

### E. Modes of modulation and conditions for mechanism-based activation

It is now possible to analyze the thermodynamic conditions on the binding of a modulator A in order to achieve mechanism-based sirtuin activation under the rapid equilibrium segments approximation, and to consider the predicted changes in the steady state, equilibrium and dissociation constants in the reaction mechanism of this enzyme.

As we have mentioned above, *v*_max_*/*[*E*]_0_ is approximately constant within this family of mechanisms if the *K*_*d,A*_’s for binding of the modulator A to the complexes in the reaction mechanism satisfy condition (iii). As long as the modulator does not greatly increase the binding affinity coproduct binding affinity and reduce the coproduct dissociation rate (for example, the stabilization of a closed loop conformation), this condition reasonable to assume. The latter effect can render product dissociation rate limiting and reduce *k*_*cat*_, as observed in the case of Ex-527^34^. If the modulator reduces *k*_*cat*_, the net extent of activation will be reduced. But note that, even if this occurs, reduction in *k*_*cat*_ will not alter catalytic efficiency *k*_*cat*_*/K*_*m*_. It is justified to omit binding of A to the coproduct complex from the mechanism-based activation model. In addition, if the rate limiting chemistry step is much slower than product release, binding of A will not significantly reduce *k*_*cat*_.

In order to achieve mechanism-based activation, it is necessary to increase *k*_*cat,app*_*/K*_*m,app*_. According to the equations, *K*_*m,NAD*_+,*app* will be smaller than *K*_*m,NAD*_+ if

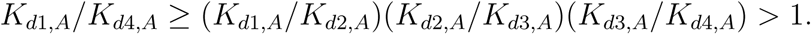

We consider one by one each of these three respective ratios of *K*_*d,A*_’s (or equivalently, the ΔΔ*G*’s of the *NAD*^+^ binding, exchange, and NAM binding reactions) induced by A binding, in order to identify mechanisms by which this can occur in terms of the steps in the enzymatic reaction.

According to the equations,*K*_*d*1,*A*_*/K*_*d*2,*A*_ *<* 1 would mean that binding of the modulator A enhances the binding affinity of the cofactor *NAD*^+^ to the E.Ac-Pr complex. Although this is the principal means by which allosteric activators increase enzyme activity, it is not the only possible means for mechanism-based activation. It is also possible in principle for a mechanism-based activator to reduce *K*_*m,NAD*_+ even if *K*_*d*1,*A*_ *≥ K*_*d*2,*A*_. In this scenario, in order for *K*_*m,NAD*_+,*app < K*_*m,NAD*_+, it must be the case that (*K*_*d*2,*A*_*/K*_*d*3,*A*_)(*K*_*d*3,*A*_*/K*_*d*4,*A*_) *> K*_*d*1,*A*_*/K*_*d*2,*A*_ or equivalently, (*K*_*d,NAM*_′*/K*_*ex*_′)(*K*_*ex*_*/K*_*d,NAM*_) *> K*_*d,NAD*+_′*/K*_*d,NAD*+_. The decrease in *K*_*m,NAD*_+ can then be caused by modulation of the exchange rate constants that results in a reduction in *K*_*ex*_, an increase in *K*_*d,NAM*_, or both of these effects. Please note that if *K*_*d,NAM*_ is altered by the presence of modulator, we have mixed noncompetitive inhibition^28^ of base exchange (Fig. 2).

As shown in our prior modeling studies of sirtuin enzymes^28^, the nicotinamide moiety of *NAD*^+^ makes very similar interactions with the enzyme prior to and following bond cleavage. The main distinction is a conserved side chain conformational change (e.g., Phe33 in Sir2Tm, Phe157 in SIRT3) which results in the destabilization of NAM binding following cleavage of the bond^35,36^. Because the binding of NAM is already destabilized in this way by the native protein conformation, and since under rapid NAM dissociation (approximation iii above, which is believed to hold for sirtuins) *k*_2_ and *k*_−2_ do not appear in the expression for *K*_*m*_, it is likely that 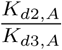 makes the dominant contribution to 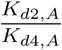 for activators. We note that our model accommodates the possibility of arbitrary combinations of ΔΔ*G*_*ex*_ and ΔΔ*G*_*bind,NAM*_, even though the value of *K*_*d*2_*/K*_*d*4_ needed for activation is likely to be achieved primarily by changing the free energy change of the nicotinamide cleavage reaction.

We assume that both *K*_*d,NAM*_ ‘s – those for NAM dissociation from the E.Ac-Pr.NAM and E.ADPR-Pr-Im.NAM complexes – are about equal in the presence of A binding. Given that A is assumed to not directly interact with the peptide or ADPR moiety, this is reasonable. We hence have:

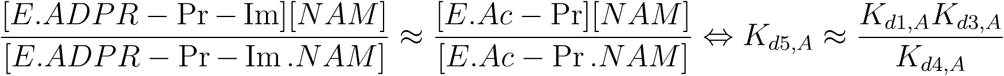

Coming back to the expression for *K*_*m,NAD*_+,*app* and substituting (1 + [*A*]*/K*_*d*2,*A*_)*/*(1 + [*A*]*/K*_*d*1,*A*_) *≥* 1, the rapid equilibrium assumptions applied to this example system imply that in order to activate the enzyme at [NAM]=0, unless A improves cofactor binding affinity, it must increase *k*_1_ (*k*_1,*app*_ *> k*_1_), *k*_2_ (*k*_2,*app*_ *> k*_2_) or both. These two scenarios cannot be distinguished under the rapid equilibrium segments model. It is unlikely that an increase in *k*_1_ will achieve activation A if increases *K*_*d,NAD*_+,_*app*_. An increase in *k*_2_ means the rate of nicotinamide cleavage is accelerated. Under the rapid equilibrium segments framework, this occurs through preferential stabilization of the E.ADPR-Pr-Im complex by the modulator. We note that nicotinamide cleavage induces structural changes (for example, unwinding of a helical segment in the flexible cofactor binding loop^37,38^) across all sirtuins studied, and such changes might enable preferential stabilization of the E.ADPR-Pr-Im complex, in a manner similar to the stabilization of specific loop conformations by mechanism-based inhibitors^34^. In fact, in crystallographic studies of reported activators of sirtuins other than SIRT3, including both long-chain fatty acids^19,31^ and recently reported activators of SIRT5 and SIRT6^18^, stabilization of alternative, non-native conformations of this loop has been observed.

Because an open loop conformation favors *NAD*^+^ binding, if the modulator stabilizes a closed loop conformation, the following thermodynamic conditions on the binding of A to the complexes in the sirtuin enzyme mechanism are expected:

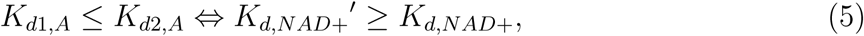

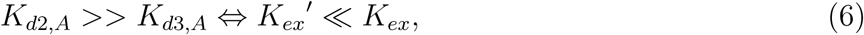

where the ≫ sign means that *K*_*d*2,*A*_*/K*_*d*3,*A*_ *> K*_*d*1,*A*_*/K*_*d*2,*A*_. We also require the necessary but not sufficient condition that *K*_*d*2,*A*_*/K*_*d*4,*A*_ *> K*_*d*1,*A*_*/K*_*d*2,*A*_. As we have discussed above, since NAM is believed to already dissociate quickly from the native active sites of sirtuins, increasing *K*_*d*3,*A*_*/K*_*d*4,*A*_ (which would destabilize NAM binding) is not considered as a mode of activation. Decrease in *K*_*ex*_ corresponds to hyperbolic (or partial) noncompetitive inhibition^28^ of base exchange/activation of nicotinamide cleavage (as opposed to complete quenching of the base exchange reaction; see Fig. 2), within the conventional nomenclature.

Fig. 10 depicts the model-predicted changes to the various steady state, Michaelis and dissociation constants in the sirtuin reaction mechanism in the presence of such a modulator. The subfigures demonstrate how varying substrate and product concentrations, respectively, provide complementary information required to elucidate the activator’s mechanism of action.

**FIG. 10.**
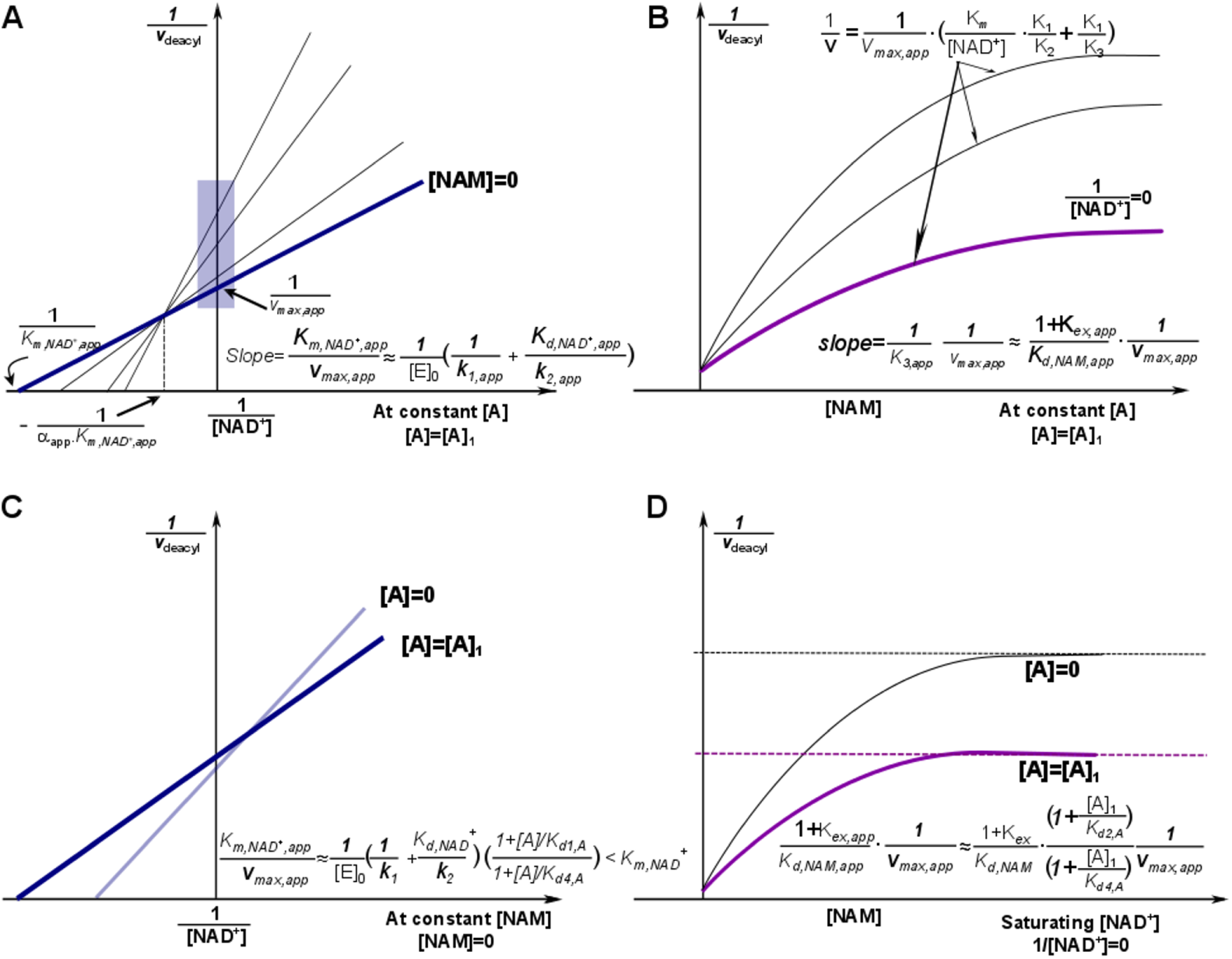
Mechanism-based activation of sirtuin enzymes: predicted steady-state properties and dose-response behavior. (A) Double reciprocal plots for deacylation initial rate measurements in the presence of activator. The blue box on the y-axis highlights the data that is used to construct the Dixon plot at saturating [NAD+] depicted in (B). (B) Dixon plots for deacylation initial rate measurements in the presence of activator. The arrows point to predicted plateaus in these curves. (C) Comparison of double reciprocal plots at [NAM] = 0 uM in the presence and absence of activator. (D) Comparison of Dixon plots at 1/[NAD+] = 0 in the presence and absence of activator. A denotes a mechanism-based sirtuin activating compound. Note that the model depicted omits the term quadratic in [NAM] in eqn (1) and the plateaus/dotted lines shown in the Dixon plots are the asymptotic values to which the model-predicted rates converge in the absence of this term.

Further discussion of model-predicted changes to the various steady state, Michaelis and dissociation constants in the sirtuin reaction mechanism in the presence of such a modulator is provided in^21,33^. We emphasize that these are properties of a lead compound for mechanism-based enzyme activation; not all of these properties must be displayed by a hit compound. In particular, the fact that the hit compound HKL requires further improvement in several of these properties to qualify as a steady state activator does not preclude it from being a non-steady state activator, as discussed further below. Note that the increasing value of *α* (intersection point of lines in double reciprocal plot closer to y axis in Fig. 8) for the p53 substrate in the presence of honokiol is consistent with the decrease in cofactor binding affinity measured by microscale thermophoresis. Moreover, the computationally predicted interaction of HKL to the flexible cofactor binding loop (Figs. 5,6), which will alter the loop conformation, is fully consistent with its steady state effects on SIRT3-catalyzed deacetylation delineated above.

Further details are provided in references^21,28^. Reference^33^ also reports the application of experimental characterization methods for mechanism-based activation, to honokiol as well as other proposed hit compounds, within the context of such a modulation model.

### F. Apparent steady state effects of unsaturating concentrations of honokiol on SIRT3

Although saturating [*A*] (as applied above for the steady state characterization of HKL) is needed to characterize the effects of a hit compound for mechanism-based activation on the chemical and binding rate constants of the reaction, the hit compound can be applied at unsaturating concentration in order to leverage the positive effects of *A* on certain steps of the reaction while reducing its negative effects on other steps. At unsaturating [*A*], the reaction can follow a path between the front and back faces of the cube resulting in apparent steady state constants as shown in Fig. 2. These effects which combine front and back face rate constants, are represented by the higher order terms in the expressions for the apparent steady state constants, such as that for *K*_*m,NAD*+,*app*_ derived above. These higher order terms are required to understand the effects of higher [*A*].

As such, note that the physiological effect of HKL activation was reported 10uM concentration^20^. Consistently it was shown above in Fig. 7 and also in reference^33^ that 10uM HKL activates SIRT3 under non-steady state conditions. By contrast, at this concentration, according to reference^33^, there is very little net effect of HKL on activity under steady state conditions. Note that 10uM HKL is above the *K*_*d*_ of the compound as reported in reference^33^. However, the effect of HKL on activity in the steady state dose response curve reported in reference^33^ extends above 100uM HKL. This is because the rate of dissociation/association of HKL from/to the enzyme with respect to the other rate constants of the reaction can affect the apparent values of the steady state rate constants in Fig. 2 and this effect is essentially eliminated at HKL concentrations above 100uM.

On the other hand, unsaturating [HKL] can allow the apparent rate constants for steps such as coproduct release to be increased with respect to the values of these rate constants in the presence of HKL (the rate constants on the back face of Fig. 2). Nonetheless, hit-to-lead evolution and lead optimization of mechanism-based enzyme activating compounds should be carried out at saturating [*A*] (as applied above) in order to increase the robustness of the hit compound with respect to physiological expression level and substrate level differences.

### G. Combining non-steady state and steady state data to characterize SIRT3 modulation by honokiol

As shown in reference^20^, the effect of HKL on SIRT3 is physiologically relevant, reducing cardiac hypertrophy in mice. Reference^20^ did not distinguish between non-steady state and steady state activation, or between saturating and nonsaturating effects of HKL. Nonetheless, the results therein combined with those in Fig. 7 above and those of^33^ show that non-steady state rate enhancement under unsaturating HKL can have physiologically favorable effects. Fig. 7 above shows between 50 − 100% activation of SIRT3 by HKL for the p53-AMC substrate and reference^33^ shows activation for the MnSOD substrate, both under non-steady state conditions. Reference^20^ established the activation of the SIRT3 activity on the MnSOD substrate under larger ratios of enzyme to substrate and enzyme to HKL concentrations, which exaggerates non-steady state effects. On the other hand, steady state rate enhancement under saturating conditions of the modulator would be expected to have more significant and robust effects.

The enhancement of the initial rate but not the steady state rate by HKL shows it is not an allosteric activator. In such a system it is important to distinguish between steady state and initial/non-steady state rates, as described in detail in^33^.

Under conditions of significant enzyme excess, the reaction follows pseudo-first order kinetics with respect to the enzyme. In our modeling of this system, we will apply the same assumptions that we have applied in our prior steady state modeling of sirtuins^21,28,33^: a) [*AADPR*] can be omitted from the state vector since its off-rate dominates its effect on the reaction rate; b) the rate of formation of Deac-Pr can be measured to follow the reaction, but [*Deac* − *Pr*] can be omitted from the state vector. The non-steady state dynamics of the system are then described by the following differential equation (under conditions of unsaturating NAD+ and saturating Ac-Pr):

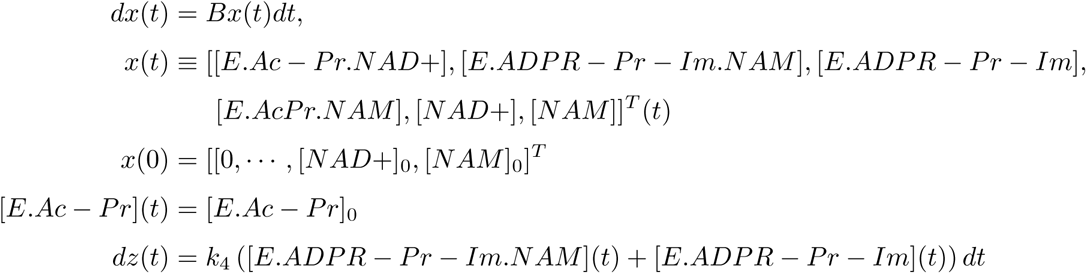

where *x*(*t*) denotes the vector of species in the reaction mechanism under pseudo-first order conditions with respect to *E.Ac* − *P r, z*(*t*) denotes the measured quantity ([*Deac* − *P r*]), and where

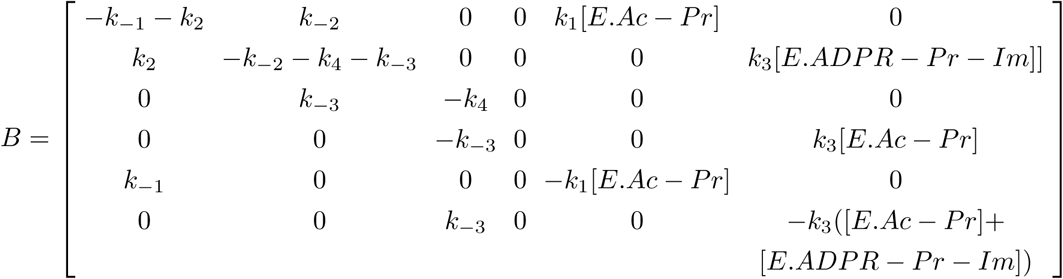

This system is quadratic (nonlinear) when [*NAM*] is low but not negligible. If [*NAM*]_0_ is sufficiently high it is linear. In addition, if we assume that like *O* − *AADPR* its on-rate is sufficiently low especially at early times, [*NAM*] can be omitted from the dynamical system, resulting in a 5 component state vector.

Note that a feature of the solution to this system, unlike the corresponding steady state system, is that a reduction in rate of regeneration of *E.AcPr* does not reduce the rate of product formation since [*E.AcPr*] is invariant, whereas in the steady state system [*E.AcPr*] equals [*E*]_0_ minus the sum of concentrations of all other reaction species. Thus comparing the rates of product formation for the steady state vs non-steady state system allows determination of whether the rate limiting step is product release or the final chemistry step of the reaction. In addition, the estimated rate constant *k*_4_ in the steady state system is the slower of the final chemistry step and product release, whereas in the non-steady state system it is the chemistry step. In the non-steady state system reaction diagram corresponding to Fig. 1, the two arrows pointing from *E.ADPR* − *Pr* − *Im* and *E.ADPR* − *Pr* − *Im.NAM* to *E.AcPr* (*E.S*_2_) no longer point to that vertex, because [*E.AcPr*] = [*E.AcPr*]_0_.

Therefore the pre-steady state activation and non-steady state activation of SIRT3 by HKL, as reported in reference^33^, indicate that HKL reduces the rate of product release (*k*_4_), but not necessarily the rate of the final chemistry step. This is fully consistent with the increase in AAPDR coproduct binding affinity in the presence of HKL reported in^33^. In Section 5 below, simulations are reported that further corroborate this result. Therefore *k*_4_ in the matrix *B* above is not the same as *k*_4_ in the steady state model in the presence of HKL, since the latter being the rate limiting step of product release and the final chemistry step, is dominated by the rate of product release. On the other hand, in either the presence or absence of HKL, *k*_4_ in the matrix B above is the rate of the second (deacetylation) chemistry step. Importantly, note that in the expression for the steady state catalytic efficiency *k*_*cat*_*/K*_*m,NAD*+_, which is relevant to steady-state sirtuin activation under NAD+ depletion conditions associated with aging, the rate constant *k*_4_ does not appear. Thus the reduction in *k*_4_ by such a modulator does not affect catalytic efficiency. Hence the non-steady state activation by HKL is relevant to the goal of increasing catalytic efficiency, but further hit-to-lead evolution is required (see below).

Measurements on the non-steady state system do not provide information on the effect of a modulator on product release, whereas measurements on the steady state system do not distinguish between the effects of the modulator on the rate of product release vs its effects on the rate of the final chemistry step. Hence the two types of measurements are complementary in the characterization of hit compounds for mechanism-based activation.

Although the non-steady state dynamical equation has an analytical solution under conditions of enzyme excess, the solution is exponential in the rate parameters. The eigenvalues of the matrix *B* must be obtained numerically. The effect of HKL on these eigenvalues determine the extent of rate enhancement under non-steady state conditions. Moreover, the eigenvalues of B can be compared to the expression for *k*_*cat*_*/K*_*m*_ to uncover the relationship between pre-steady state activation and catalytic efficiency. In order to identify the effects of HKL on these parameters that result in activation under non-steady state but not steady state conditions, the methods described in reference^29^, which achieve system identification through a combination of steady state and/or linear Kalman filtering analyses, can be applied. Note that reference^29^ focused on system identification for directed evolution of enzymes and did not account for modulation of enzymatic activity by small molecules, the focus of the present work.

## V. SIMULATIONS: SIRTUIN ENZYMES

In this Section, we present simulation results applying the structural theory of mechanism-based activation presented in Section 2 to sirtuin enzymes.

### A. Simulation methods

#### Loop modeling

The following protocol was used to prepare starting structures for molecular dynamics simulation of the SIRT3 ternary (SIRT3+AcPr+NAD+), intermediate (SIRT3+ADPR-Pr-Im) and product (SIRT3+ Pr +OAADPR) complexes with native and nonnative loop. For the ternary complex (prepared based on the 4FVT crystal structure for all protein residues except the cofactor binding loop), the intermediate (closed) conformation of the cofactor binding loop (residues 155-178) was substituted, whereas for the intermediate complex (prepared based on the 4BVG crystal structure), the ternary (open) conformation of the cofactor binding loop (residues 155-178) was substituted. For the coproduct complex (prepared based on 4BVH crystal structure), the apo (open) conformation of the cofactor binding loop (residues 155-178 from the 3GLS crystal structure) was substituted. The following steps were applied:

1. Sequence-structure based superimposition
2. Graft the coordinates for the co-factor loop region.
3. Run protein preparation wizard on the modelled product complex (open/closed loop conformation).
4. Predict/repack the side chains for all residues within 7.5A of the grafted loop region using Monte Carlo approach together with backbone sampling.
5. Carry out prime energy refinement only on those residues which were repacked keeping the others fixed.

#### Side chain prediction

Rotamer-based side chain prediction was applied following loop replacement. A subset of residues was chosen for side chain prediction. The residues considered for prediction are 144-180,195,199,204,207,210,227-234,248,251,291,294,324 – residues with 7.5A of a modelled loop region were refined. Native side chain conformations were repredicted in native environment prior to prediction in non-native environments.

Four methods for side chain prediction in the protein local optimization program (PLOP) were used

1. Default method – No backbone sampling or reorientation of the CA-CB bond is performed.
2. Monte Carlo approach – Monte-Carlo sampling of side-chain conformations including backbone flexibility.
3. CA-CB vector sampling – varying the orientation of the CA-CB bond by up to 30 degrees from the initial direction.
4. Backbone sampling – Sample the backbone on a set of 3 residues centered on the residue for which the side chain is being refined.

We used the top 3 conformations provided by the Monte Carlo run for minimization.

#### Molecular dynamics simulation

- **Force field**. The Amber99SB force field^39,40^ was used for all the molecular mechanics calculations. Extra parameters were adapted from the following sources: parameters for Zn developed by Luo’s group^41^; parameters for NAD+ developed by Walker et al^42^ and Pavelites et al^43^; parameters for acetylated lysine developed by Papamokos et al^44^. Amber14 tools (Antechamber program) are employed for parametrizing the non-standard residues and tleap program for creating topology and force field parameters for the standard residues.
- **Fast long-range electrostatics**. The Particle Mesh Ewald (PME) method^45^ was used to calculate the electrostatic energy.
- **Explicit vs implicit solvation**. Explicit solvent for sampling but implicit was used for more energy calculations Water molecules were stripped from the molecular dynamics trajectories to prepare for MM/PBSA and MM/GBSA re-runs.
- **MD simulation**. All molecular dynamics simulations were performed with periodic boundary conditions to produce isothermal-isobaric ensembles (NPT) at 300 K using the NAMD program^46^. The integration of the equations of motion was conducted at a time step of 2 femtoseconds. The covalent bonds involving hydrogen atoms were frozen with the SHAKE algorithm^47^. Temperature was regulated using the Langevin dynamics with the collision frequency of 1 ps-1. Pressure regulation was achieved with isotropic position scaling and the pressure relaxation time was set to 1.0 picosecond.
- **Equilibration vs production simulations**. There were three phases in the molecular dynamics simulations. First, in the relaxation phase, the system underwent a 2000-step minimization before a short 200 ps NPT molecular dynamics simulation, with the main chain atoms of the protein restrained to the positions in the crystal structures with force constants of 5 kcal mol-1 Ã-2. Next, the systems ran for various lengths of time up to 22 ns in the equilibration phase. Last, the sampling phase included a 10 ns of molecular dynamics simulation. molecular dynamics (first stage): Before launching production run we minimize and equilibrate (200 ps of equilibration) the system and check for the stability (time vs RMSD, time vs potential energy, time vs pressure, time vs temp) to ensure that the system is stable before launching the production run (15 ns). This entails a slow heating up (from 15 k to 300 K with a gradual increase of 15 K) of the system and then running equilibration simulation for 200 ps at 300K.

#### Estimation of conformational and binding energies

Energies were calculated using the MM-PBSA and the MM-GBSA methods as implemented in the AMBER package^48^. In MM-PBSA and MM-GBSA, binding free energy is evaluated as:

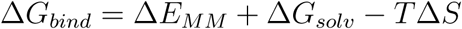

where = Δ*E*_*MM*_, + Δ*G*_*solv*_ and *T* Δ*S* are the changes of gas-phase interaction energy, solvation free energy, and solute conformational entropy change upon binding, respectively. Δ*E*_*MM*_ includes internal energy in bonded terms, electrostatic and van der Waals energies. Δ*G*_*solv*_ is the sum of polar contributions calculated using the PB or GB model, and nonpolar contributions estimated from solvent-accessible surface area (SASA). All the calculations were carried out using the MMPBSA.py module with AmberTools13^48^. The polar contribution of the solvation free energy was calculated by the GB model developed by Onufriev et al.^49^ and by the PB method implemented in the pbsa program. A salt concentration of 0.1 M was used in MM-GBSA calculations. The solvent-accessible surface area was evaluated using the LCPO method^50^. Because relative free energy trends were of interest, solute conformational entropy change was neglected. Energies were evaluated using 10000 snapshots extracted from the last 10 ns at a time interval of 1 ps for each trajectory after ensuring that each one of these trajectories was completely stable.

### B. Simulation Results

Across all sirtuins studied, nicotinamide cleavage induces structural changes (for example, unwinding of a helical segment in the flexible cofactor binding loop (Fig. 11 (top)). Comparison of the crystal structure of the human SIRT3/Ac-Pr/Carba NAD ternary complex (4FVT) with SIRT3 apo structure (3GLS) shows the cofactor binding loop adopts an open conformation that is very similar in both, whereas the flexible loop in the SIRT3/ADPR-Pr-Im complex (4BVG) has a significantly higher RMSD with respect to that in 4FVT (Fig. 11 (bottom)). Stabilization of alternative, non-native conformations of this loop have been observed crystallographically by reported activators of sirtuins other than SIRT3, including both long-chain fatty acids and recently reported activators of SIRT5 and SIRT6. As shown above in Fig. 5, mechanism-based sirtuin activators like HKL directly interact with this loop, altering its conformation and affecting the rate of catalysis. As described in Section 2, the potential of mean force *ϕ*_*A*_(*R*) generated by this interaction causes free energy changes in ligand binding ΔΔ*G*_*bind,L*_ according to equation (2).

**FIG. 11.**
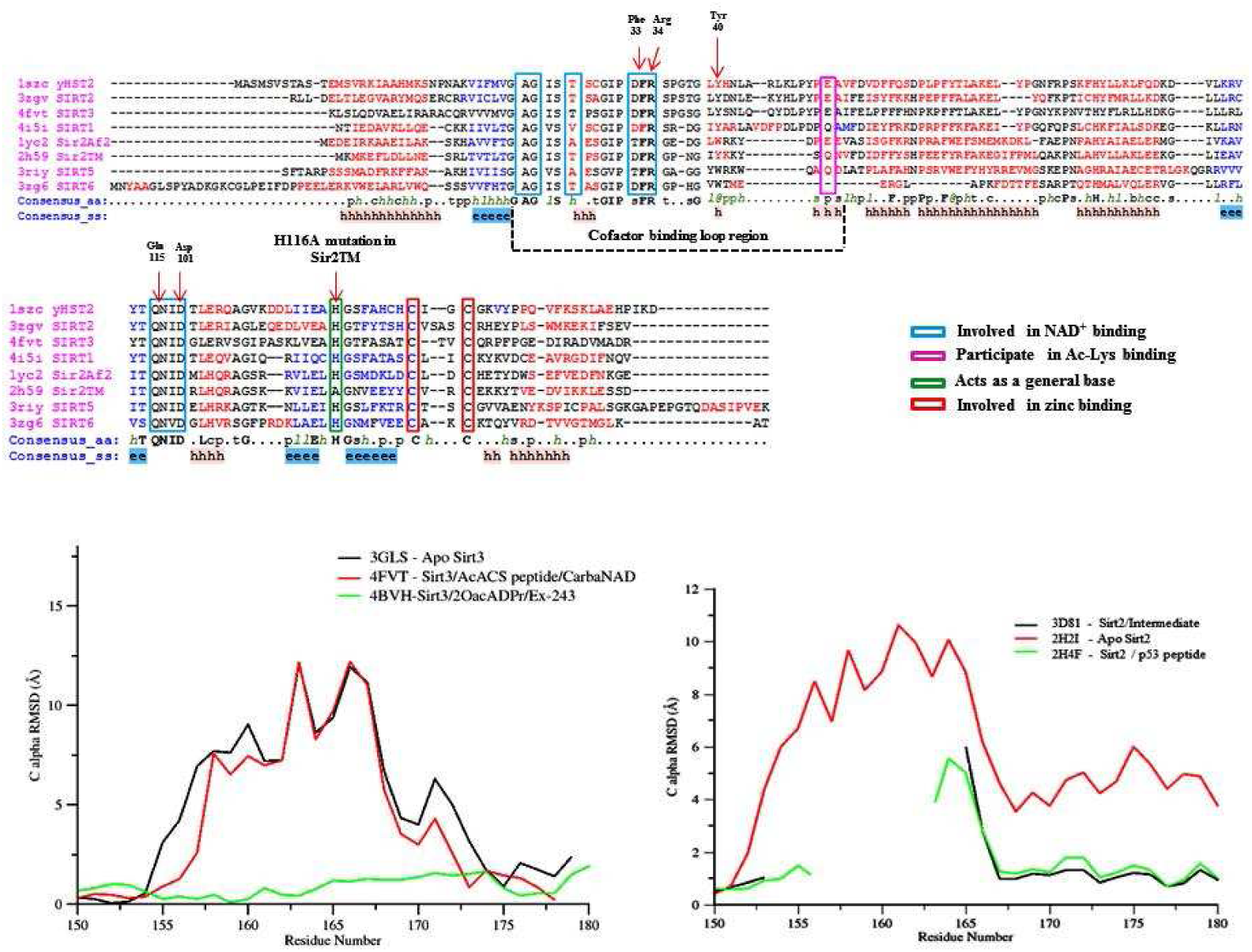
**Top**) PROMALS3D based sequence alignment of sirtuin proteins. Residues shown in the alignment are colored according to their predicted secondary structure elements (red: alpha-helix, blue: beta-strand). The boundaries of the co-factor binding loop region are highlighted using black dotted lines. The consensus sequence (consensus aa) and consensus predicted secondary structure (consensus aa) are shown at the bottom of the alignment. Consensus amino acid symbols are represented by: conserved amino acids are in bold and uppercase letters; aliphatic: l; aromatic: at; hydrophobic: h; alcohol: o; polar residues: p; tiny: t; small: s; bulky residues: b; positively charged: +; negatively charged: -; charged: c. The global consensus predicted secondary structure are represented by alpha helix (h) and beta strand (e). Residues important for co-factor binding, substrate binding and catalysis are highlighted in colored boxes. **Bottom**) Shown in the left panel are Sirt3 proteins and their per-residue RMSD values for the cofactor binding loop region computed over all atoms with reference to crystal structure of a Sirt3 intermediate complex (4BVG). The right panel shows RMSD values for Sir2Tm proteins calculated with reference to crystal structure of a Sir2 ternary complex (2H59). Residues (155-178) correspond to the co-factor binding loop region. Unresolved loop regions are not plotted in the figure.

As such, in the present study molecular dynamics simulations have been carried out on the SIRT3/Ac-Pr/NAD+, SIRT3/INT(/NAM) and SIRT3/Pr/AADPR complexes in the catalytic mechanism (Fig. 12). Local side chain degrees of freedom are optimized under a molecular mechanics energy function and QM partial charges were derived and fit for all ligands. Modern loop prediction methods have improved energy function accuracy and enhanced sampling, but sampling and energy errors are sometimes encountered for very long loops. Loop replacement required minimal backbone sampling, and both energy function and sampling accuracy was validated for side chain optimization, which produced sub-angstrom RMSDs in prediction of native complex side chain packing.

**FIG. 12.**
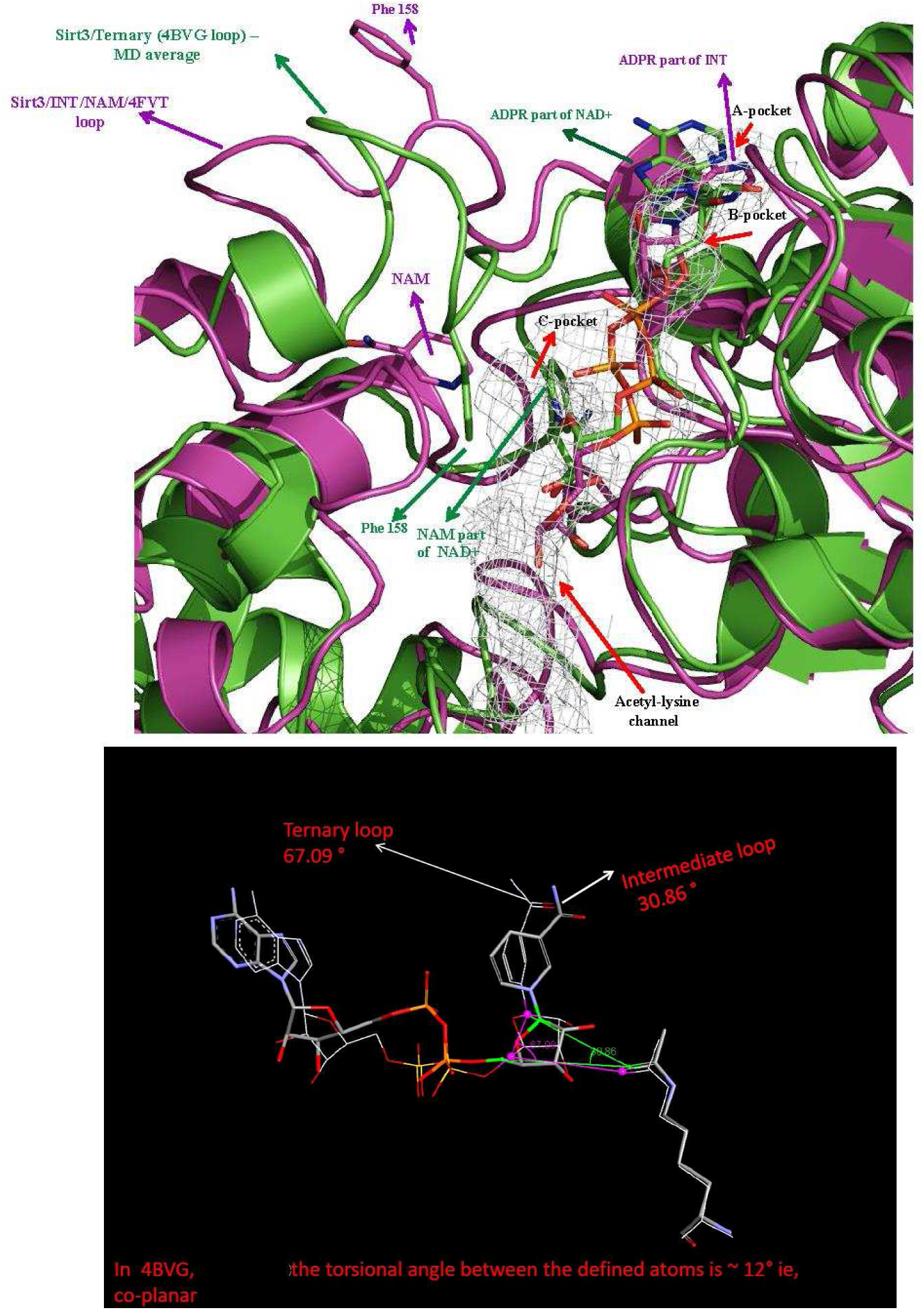
**Top**) Superposition of (purple) the time averaged molecular dynamics structures of Sirt3/ADPR/NAM intermediate complex modeled based on the crystal structure of intermediate complex (4BVG) with the co-factor binding loop residues (155-178) being replaced with those from the ternary complex crystal structure (4FVT) and (green) the Sirt3/AcPr/NAD+ ternary complex modeled based on the crystal structure of ternary complex (4FVT) with the co-factor binding loop residues (155-178) being replaced with those from the intermediate complex crystal structure (4BVG). Differences in the conformations of the co-factor binding loop and the position of the Phe residue and NAM are highlighted. Individual subsites are highlighted. **Bottom**) Close-up of NAD+ in the ternary complex for open vs closed loops showing order parameters from molecular dynamics averages.

These simulation results thus consider two dominant backbone conformations, which constitute the local DOF (*r*_1_, …, *r*_*l*_) in eqn 1. It is assumed that the closed loop conformation in the ternary complex corresponds to *R*_0_ and the potential of mean force *ϕ*_*A*_ destabilizes conformations with nonzero 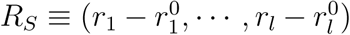. In subsequent work, we will report the values of ΔΔ*G* induced by such a potential of mean force according to eqn 2.

Free energies of binding were estimated for each of the major complexes in the sirtuin reaction mechanism, with the open and closed loop conformations of the cofactor binding loop. The results are provided in Table 1. From these calculations it is possible to estimate the changes in the Δ*G* of steps in the catalytic mechanism induced by the modulator. Further insights can be obtained by carrying out analogous calculations for the transition states as modeled by mixed quantum/molecular mechanics (QM/MM) methods, which will be employed in future work.

**Table 1:** Ligand binding affinities in sirtuin (SIRT3) complexes and effect of loop conformation.

| Complex (loop conformation) — | NAD+ | Ac-Pr / | ADPR-Pr-Im | NAM / | O-AADPR | Deac-Pr |
| --- | --- | --- | --- | --- | --- | --- |
| Panel A: MM-GBSA |  |  |  |  |  |  |
| NAD+, Ac-Pr (open) | −62.26 | −52.22 | - | - | - | - |
| NAD+, Ac-Pr (closed) | −56.23 | −46.97 | - | - | - | - |
| ADPR-Pr-Im, NAM (open) | - | - | −104.12 | −20.33 | - | - |
| ADPR-Pr-Im, NAM (closed) | - | - | −124.92 | −22.50 | - | - |
| AADPR, Deac-Pr (open) | - | - | - | - | −85.08 | −99.79 |
| AADPR, Deac-Pr (closed) | - | - | - | - | −89.60 | −19.24 |
| Panel B: MM-PBSA |  |  |  |  |  |  |
| NAD+, Ac-Pr (open) | 3.41 | −12.51 | - | - | - | - |
| NAD+, Ac-Pr (closed) | 14.57 | −15.22 | - | - | - | - |
| ADPR-Pr-Im, NAM (open) | - | - | −24.51 | −3.96 | - | - |
| ADPR-Pr-Im, NAM (closed) | - | - | −27.95 | −7.73 | - | - |
| AADPR, Deac-Pr (open) | - | - | - | - | −8.54 | −6.89 |
| AADPR, Deac-Pr (closed) | - | - | - | - | −35.94 | 3.70 |

Whereas the *NAD*+ binding affinity decreases slightly (Table 1) under the closed loop conformation (as also observed from the experimental data above on the effect of HKL on *K*_*d,NAD*+_), the acetylated lysine residue is nonetheless retained in a near-attack configuration such that *k*_2_ (which plays an important role in determining *K*_*m,NAD*+_ as discussed above), is not expected to greatly decrease. The time series of the interatomic distance between the ribose C-N carbon and acetyl oxygen is depicted in Fig. 13 for the two different loop conformations, showing that the closed loop conformation reduces the average interatomic distance and standard deviation of this distance, which is conducive to a higher value of *k*_2_. The significant increase in binding affinity of AADPR coproduct (Table 1) in the closed loop conformation compared to the open loop conformation – combined with the fact that opening of this loop is a prerequisite for AADPR release and recycling of active enzyme – is consistent with the reported increase in coproduct binding affinity in the presence of HKL^33^ and the fact that HKL that reduces the rate of AADPR release^33^. Note that AADPR binding affinity increases moreso than that of the ADPR-Pr-Im intermediate upon conformational change from the open to the closed loop conformation. However, as discussed above AADPR binding affinity does not affect the catalytic efficiency of sirtuins with respect to the [NAD+] depletion conditions relevant to aging, and has a reduced effect away from steady state under these conditions as well. In addition, the increase in binding affinity of NAM in the intermediate complex in the presence of the closed loop conformation is consistent with the experimental data reported in^33^, though the effect of this may not be very significant under low [*NAM*] conditions. Therefore, taken together the experimental and simulation data are consistent with the modulation of active site conformational degrees of freedom by HKL to enhance catalytic properties of SIRT3 under physiologically relevant conditions.

**FIG. 13.**
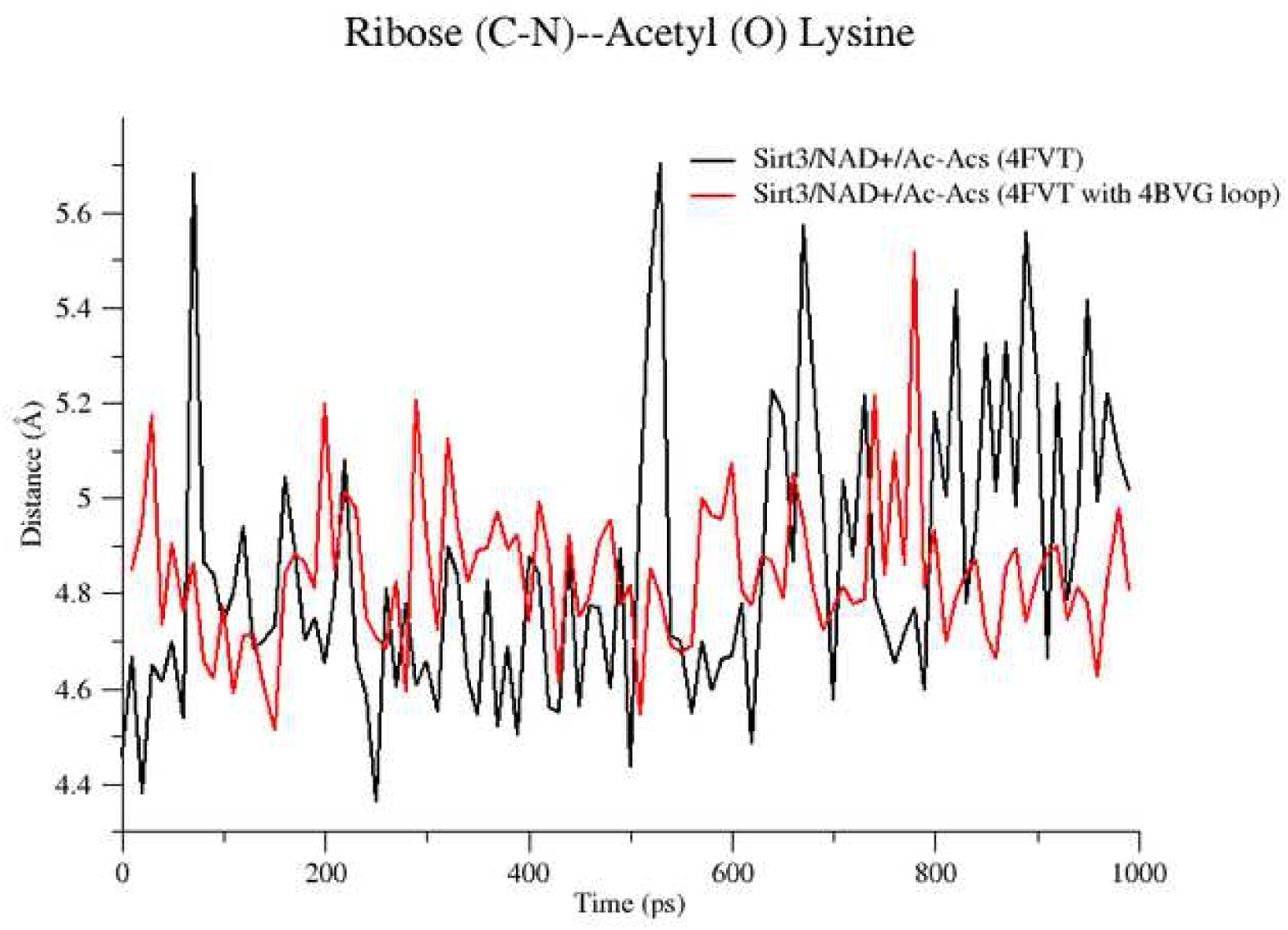
Time dependent change in distance between ribose C-N carbon and acetyl oxygen (distance plot) for 4FVT with NAD+/Ac-peptide with 4BVG loop using 12ns MD trajectory data.

In a design context, alternate dominant conformations of local degrees of freedom, such as the open and closed flexible cofactor loop conformations in the case of sirtuins, can be used together with computational methods to predict the free energy changes corresponding to the relevant rate constants. For example, the effects of loop conformational changes induced by a modulator on the binding affinity of NAD+ cofactor and the free energy change of nicotinamide cleavage can be calculated.

Indeed, a major advantage of mechanism-based activators is their scope for rational design. By contrast, design of allosteric activators for enzymes in which sites have not naturally evolved is much more challenging. Paradigms for the design of mechanism-based activators will be described in detail in a subsequent work. Briefly, one approach to designing a modulator that induces the desired free energy differences is hence to identify a potential of mean force under which the Δ*G*’s of the various reaction steps take on values that are expected to increase the steady state activity (for example, a potential of mean force that is locally harmonic around the closed loop backbone coordinates and independent of ligand coordinates), and then to match the target potential of mean force as closely as possible with a modulator’s potential of mean force (for example, by matching the force constants of the target potential of mean force to those of a second order expansion of the modulator potential of mean force around the desired backbone conformation).

Alternatively, if one does not know in advance a modulator potential of mean force that will induce the desired improvements in catalytic activity, the free energy differences in the catalytic mechanism can be calculated for trial modulator Hamiltonians using the potential of mean forces induced by the modulator Hamiltonians, in the context of a design algorithm. For example, in the case of sirtuins, one can apply an algorithm that samples molecular mutations to the hit compound in order to reduce *K*_*m*_ independently of *K*_*d,NAD*+_ and *k*_*cat*_. Assuming the observable quantity one wants to improve is catalytic efficiency, and the sampling is done with a Monte Carlo algorithm, we have the following expression for the evolutionary expectation of the catalytic activity in terms of the various free energies.

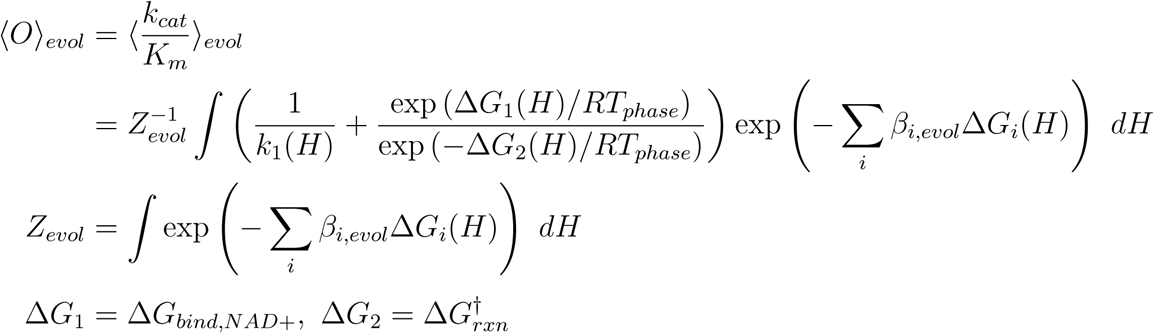

where *H* denotes the Hamiltonian of *A* and the Δ*G*’s are calculated according to the potential of mean force expressions above († denotes activation energy, here for the ADP-ribosylation reaction). Each *β*_*i,evol*_ is an inverse evolutionary temperature that governs the stringency of selection for that property (free energy difference). Then the evolutionary heat capacities, e.g.

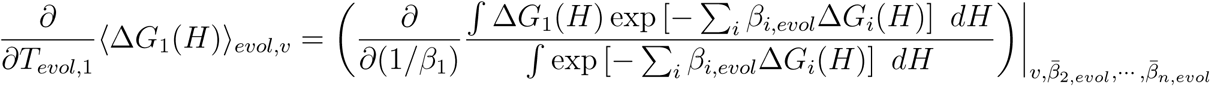

where *v* denotes evolutionary heat capacity at constant volume (meaning that only active site degrees of freedom are sampled by Monte Carlo with other degrees of freedom frozen), determine how quickly a particular free energy difference can be improved by more stringent Hamiltonian sampling. Correlations between experimentally identified and computationally predicted free energy differences can be used to calibrate the predictions. In addition, quantities that are difficult to predict through molecular modeling (such as *k*_1_(*H*) above) can be predicted using quantitative structure-activity relations (QSAR) based on experimental data.

## VI. DISCUSSION

In summary, both atomistic free energy-based and kinetic theories of mechanism-based enzyme activation have been presented in this paper. These theories have been distinguished in detail from previously reported theories of enzyme activation, including allosteric activation and activation by derepression of product inhibition. The theories were shown to be consistent with experimental data on the effects of a modulator the human SIRT3 enzyme that increases the rate of catalysis under non-steady state conditions, and simulation results on SIRT3 were presented that are also consistent with the theories. We note that the binding affinities of *A* to the transition states of the reaction were not included in the kinetic model for mechanism-based activation above; these binding affinities are never considered as part of traditional modulation models.

In future work, the free energy modulation theory introduced herein (Section 2) could be applied to co-crystallized complexes of (other) mechanism-based sirtuin activators reported in the literature, determining the potentials of mean force induced by the activator through molecular dynamics simulations. Also, the kinetic theory introduced herein (applied in Section 4 to honokiol) could be applied to kinetic data on these mechanism-based sirtuin activators in order to delineate how the compounds alter the fundamental rate constants in the reaction mechanism to increase catalytic efficiency. The modes of action of these compounds have thus far not been successfully elucidated through conventional enzyme activation theory.

## VII. APPENDIX

Derivation of expression for *K*_*mS,app*_:

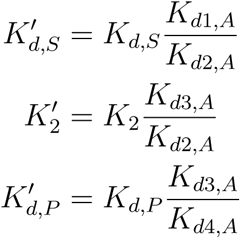

where the *K*_*d,A*_’s are the dissociation constants for A depicted in Fig. 2.

With this notation,

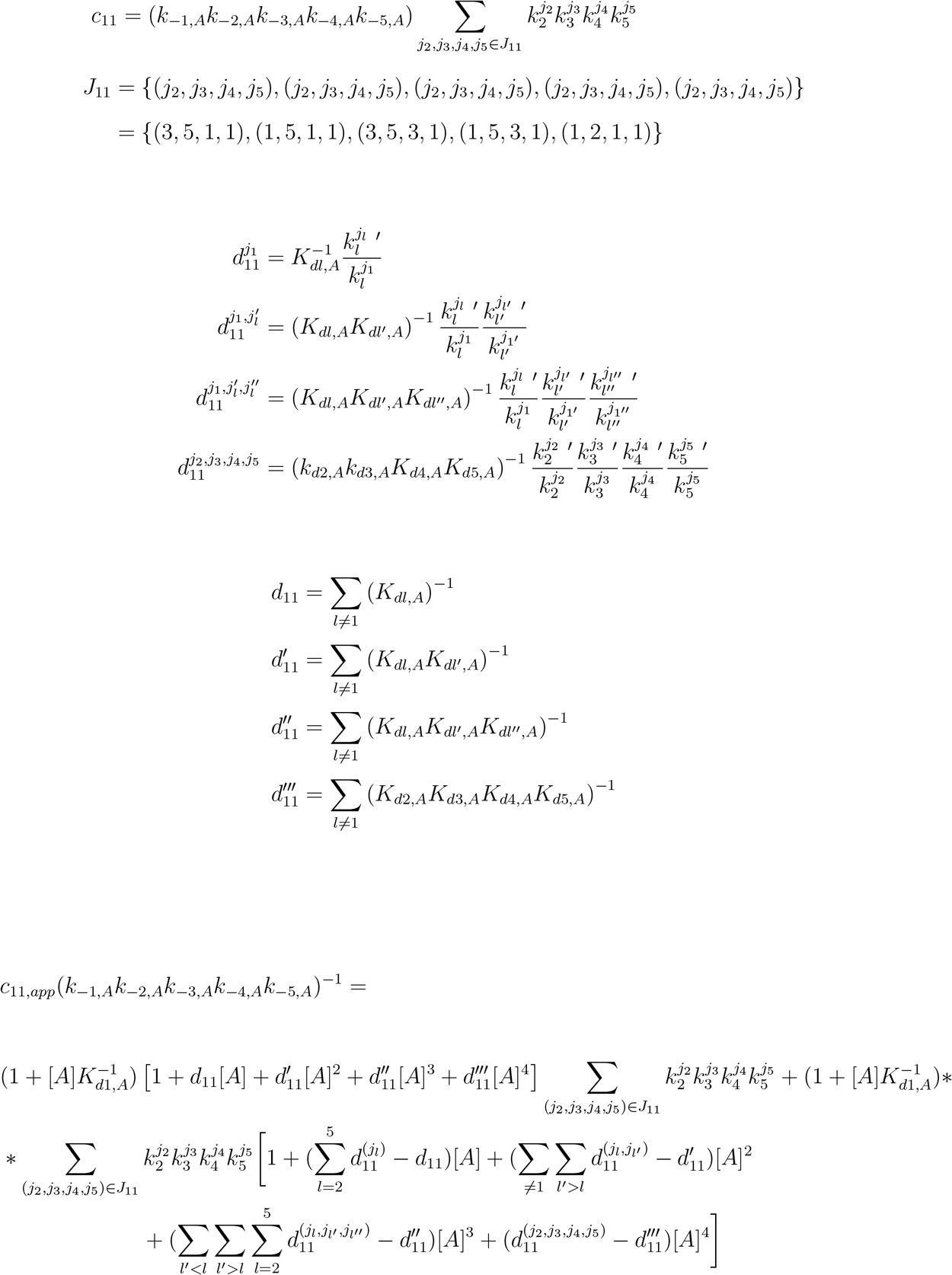

For example, in the representative subdiagrams in Fig. 3, we have

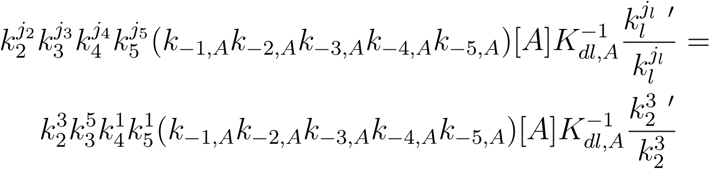

and

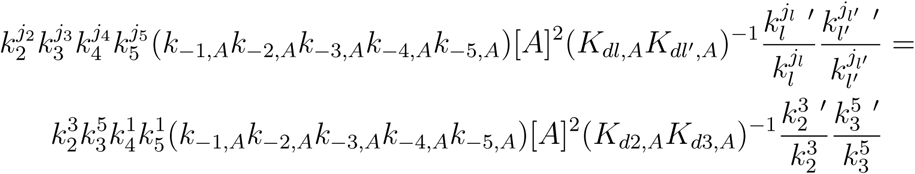

Defining

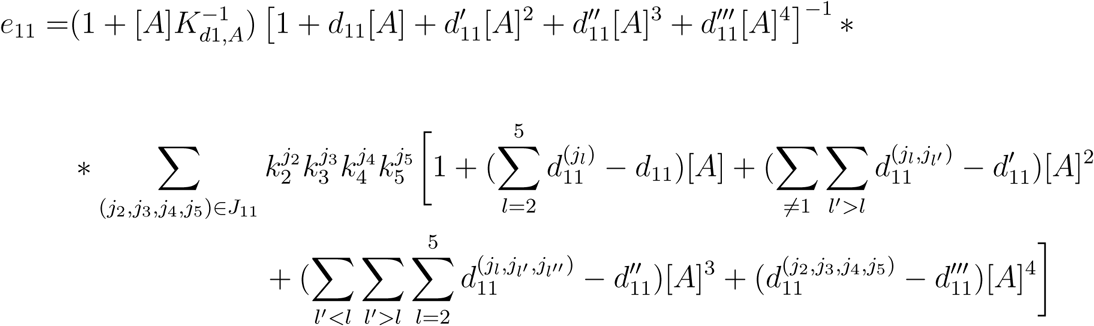

we can rewrite the expression for *c*_11,*app*_ as

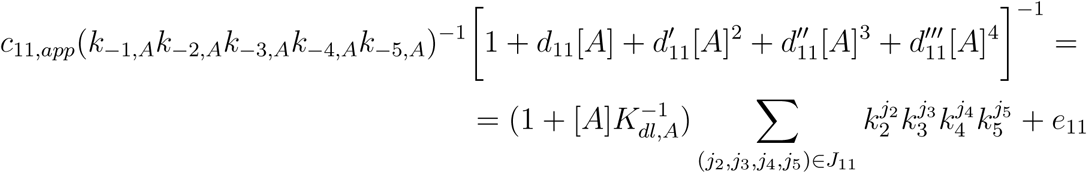

Definitions of *c*_*ij*_ for sirtuin enzymes and expressions for steady-state constants in terms of them:

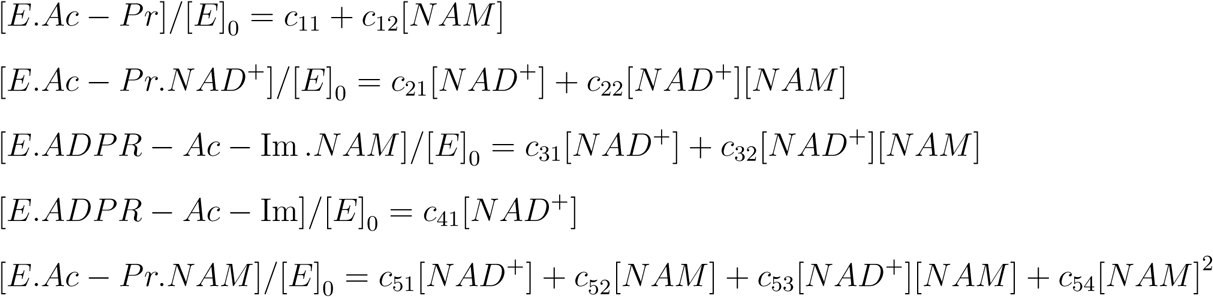

With these, the expressions for the steady state constants for sirtuin enzymes in equation (4) follow from

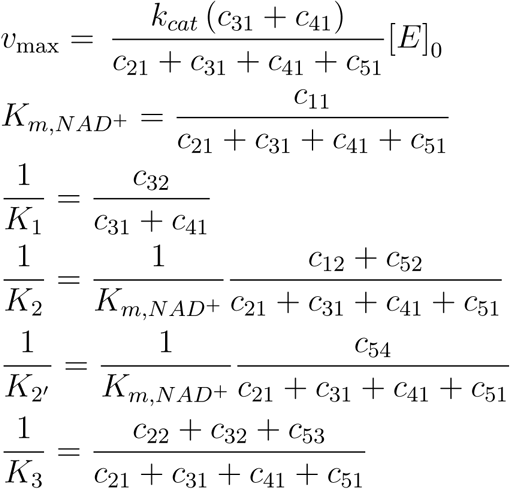

## VIII. SUPPORTING INFORMATION

See supporting information for experimental methods, supporting derivations pertaining to the modeling of mechanism-based enzyme activation, and analysis of the distinctions between mechanism-based enzyme activation and derepression of product inhibition.

## 1 Experimental methods

- **Chemicals and Reagents**. FdL2 (QPKK,AC-AMC) peptide, also called p53-AMC peptide, was purchased from Enzo Life Sciences (Farmingdale, NY). carba-NAD was synthesized by Dalton Pharma (Toronto, ON). Honokiol (HKL) was purchased from Sigma (St. Louis, MO). The solubility profiles of these compounds were determined using HPLC. All other chemicals, solvents used were of the highest purity commercially available and were purchased from either Sigma (St. Louis, MO) or Fisher Scientific (Pittsburgh, PA) or from VWR International (Bridgeport, NJ). The plasmid containing human Sirt3,102-399 (pEX-His-hSIRT3,102-399) was purchased from OriGene (Rockville, molecular dynamics).
- **Expression and purification of hSirt3**,**102-399**. The hSirt3,102-399 was expressed in E.coli Arctic Express (DE3) cells (Agilent Technologies, Wilmington, DE) as per manufacturer’s recommendations. A single bacterial colony was inoculated in LB media containing ampicillin and gentamycin at 37 degrees C. For protein purification, we inoculated overnight grown culture into 200 ml LB medium and grown at 30 degrees C, 250 rpm for 4 hours. We induced Sirt3 expression by adding 1 mM IPTG at 15 degrees C. After 24 hours, cells were harvested, and the pellet was re-suspended in A1 buffer (50 mM NaH2PO4, 300 mM NaCl, 10 mM imidazole, pH 8.0) and sonicated. A clear supernatant was obtained by centrifugation at 14000 x g for 25 min at 4 degrees C then loaded onto a 5 ml HisTrap HP column, pre-equilibrated with A1 buffer, attached to an AKTA pure FPLC system (GE Healthcare, Pittsburgh, PA). The column was then washed with 50 ml each buffer A1, A2 (50 mM NaH2PO4, 300 mM NaCl, 75 mM imidazole, pH 8.0), A3 (20 mM Tris-HCl, 2M urea, pH 6.8), and again with buffer A2. Protein was eluted with buffer B1 (50 mM NaH2PO4, 300 mM NaCl, 300 mM imidazole, pH 8.0). The protein fractions were pooled, dialyzed against dialysis buffer (25 mM Tris, 100 mM NaCl, 5 mM DTT, 10% glycerol, pH 7.5) at 4 degrees C overnight. The purity of final protein was *>* 85% as assessed by SDS-PAGE.
- **hSirt3**,**102-399-ligand binding assay by Microscale Thermophoresis**. DY-647P1-NHS-Ester conjugated hSirt3102-399 was used for the Microscale Thermophoresis (MST) studies. Briefly, the protein conjugation was done in 1x PBS pH 7.4, 0.05% Pluronic F-127, and the labeled protein was buffer exchanged into 47 mM Tris-HCl pH 8.0, 129 mM NaCl, 2.5 mM KCl, 0.94 mM MgCl2, 5% DMSO, 0.05% Tween-20. We used 2 nM labeled hSirt3 and titrated with varying concentrations of the modulators in the absence and presence of various concentrations of substrates, products and intermediates. The thermophoresis was measured (excitation/emission 653/672 nm) using a Monolith NT. 115 Pico (NanoTemper Technologies) at 25 degrees C. Dissociation constants (Kd) were determined using GraFit7 (Erithacus Software) by nonlinear fitting.
- **Effect of Honokiol on hSirt3, 102-399 deacetylation activity** Enzymatic reactions included either 2.5 mM NAD+ and 6.25 *µ*M peptide or 50 *µ*M NAD+ and 600 *µ*M peptide substrate in presence of HKL (range 0-200 *µ*M), in a buffer (50mM TRIS-HCl, 137mM NaCl, 2.7mM KCl, and 1mM MgCl2, pH 8.0 and 5% DMSO). We started the reactions by adding 5U hSirt3102-399 and incubated at 37oC for 30 minutes. We terminated the reactions by TFA.
- **hSirt3**,**102-399 kinetic analysis using Fluorolabeled Peptide**. We determined the steady state kinetic parameters of hSirt3,102-399 deacetylation using FdL2 peptide. The enzymatic reactions were carried out similar to as described above. For initial velocity determination, we performed the reaction at different [NAD+] and in presence of 100 *µ*M FdL2 peptide. We terminated the reactions at specified time by adding the 1X developer and measured the fluorescence on TECAN microplate reader. The raw data were fitted to the defined model equations using GraphPad Prism (GraphPad Software, Inc, CA).

## 2 Regimes of Interest

We apply the following representation to arrive at the relevant approximation (2) in the Section on Regimes of Interest in the main text.

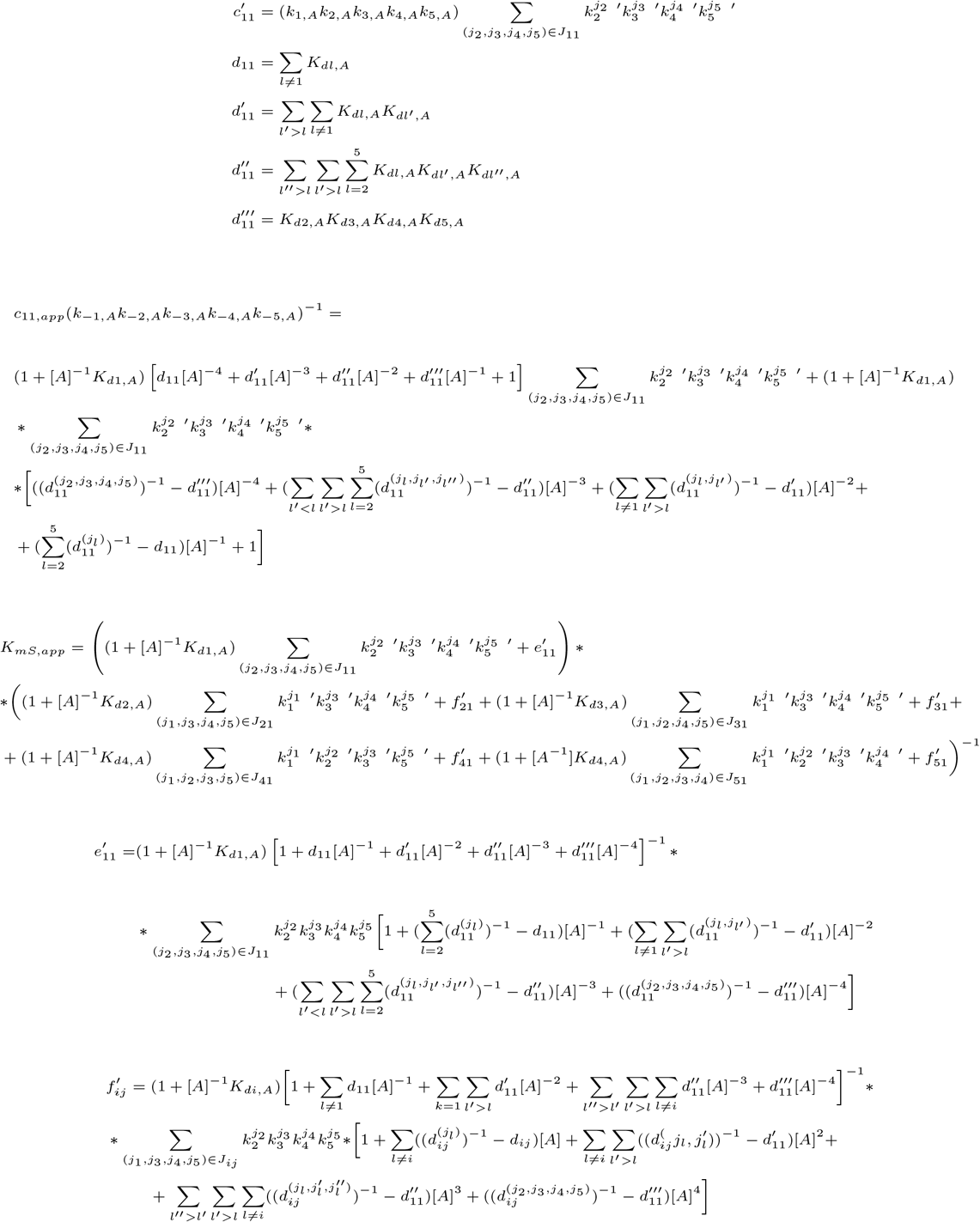

## 3 Competitive derepression of product inhibition

Generally, derepression of inhibition constitutes the only extant theory for mechanism-based enhancement of enzymatic activity, and rapid equilibrium models for such derepression have been proposed in the literature [1].

Prior work [1] on mechanism-based activation considered the use of competitive inhibitors of the reverse reaction(like isonicotinamide, isoNAM) to activate enzymes (e.g., sirtuins) at nonzero product concentration [P]. Aside from allosteric activation, this is the only other previously proposed mechanism of enzyme activation that has been experimentally investigated. These modulators rely on a favorable balance between competitive inhibition of the reverse and forward reactions for activation. Importantly, this approach cannot reduce *K*_*m,S*_. At [*P*] = 0, it will always increase the apparent value of *K*_*m,S*_ (through an increase in *K*_*d,S*_). Competitive inhibition of the reverse reaction can only reduce the *K*_*m,S*_ at nonzero [*P*].

Traditional models of depression of inhibition were not formulated for an endogenous inhibitor. The theory is significantly more complicated for an endogenous inhibitor and has not been described in the literature. Here there are two additional features: a) direct competition with the part of the substrate corresponding to the endogenous inhibitor (correlation between the effects on substrate and inhibitor binding), and b) indirect competition stemming from a reversible reaction.

Endogenous product inhibition may offer an opportunity for mechanism-based enzyme activation at [*I*] = 0, but not through previously proposed modes of action. The appropriate theory has not been previously described. Instead of competition with respect to endogenous inhibitor for binding as in the case of derepression strategies [1], then, selective stabilization of the reaction intermediate through preferential binding to structural features unique to the intermediate in particular, an altered flexible local active site degrees of freedom may be capable of activating the enzyme.

Steady state models are essential for quantitative analysis of such derepression modalities, with previously reported formulations being approximations. An extended steady state model including the small molecule modulator is required for proper analysis, given that the modulator competes with S and P to form new species rather than preferentially stabilizing certain species in the reaction mechanism. For two step enzymatic reactions with the first chemical step being reversible (of which sirtuin enzymes are an example), the figure below (Supporting Fig. 1) depicts a reaction diagram that can be used for the development of such a kinetic model, based on the unmodulated reaction diagram in Fig. 1 in the main text. The scope (and limitations) of competitive inhibition of the reverse reaction as a means of activation can be interrogated in terms of this kinetic model. Following the development of the general mechanism-based activation theory below, we describe quantitatively how the case of competitive derepression of product inhibition violates some important characteristics of mechanism-based activation. To summarize the salient issues, competitive inhibitors of base exchange will simultaneously increase the apparent value of *K*_*d,S*_ to an extent that depends on the ratio of mixed inhibition steady state constants *K*_3_ and *K*_2_. The greater the value of for an enzyme, the greater the increase in *K*_*d,S*_ that accompanies a given reduction in reverse reaction inhibition. This means that for most enzymes, a competitive inhibitor of the reverse reaction will display a significant extent of competitive inhibition of the forward reaction, at concentrations required for activation. This will reduce the maximum possible extent of activation.

### 3.1 Derepression modalities violate approximations in text

As noted above, competitive derepression of product inhibition (as reported for isoNAM in the case of sirtuin enzymes, for example) can be compared to the formalism above. In this case, the error terms will be large for all *c*_*ij*_ because the on rates will be small or off rates will be large for two of the *i*. The diagrams/terms that involve bottom face arrows will introduce these large errors. This demonstrates that the rapid equilibrium approximation is not accurate for competitive depressors of product inhibition and the on/off rates of the modulator are important. Note that *k*_*off*_ s for missing *i*s will be very large, and *k*_*on*_[*A*]s will be very small This renders it difficult for the mechanism of action of such compounds and their effects on steady state activity to be studied even approximately using binding affinities alone.

**Figure 1:**
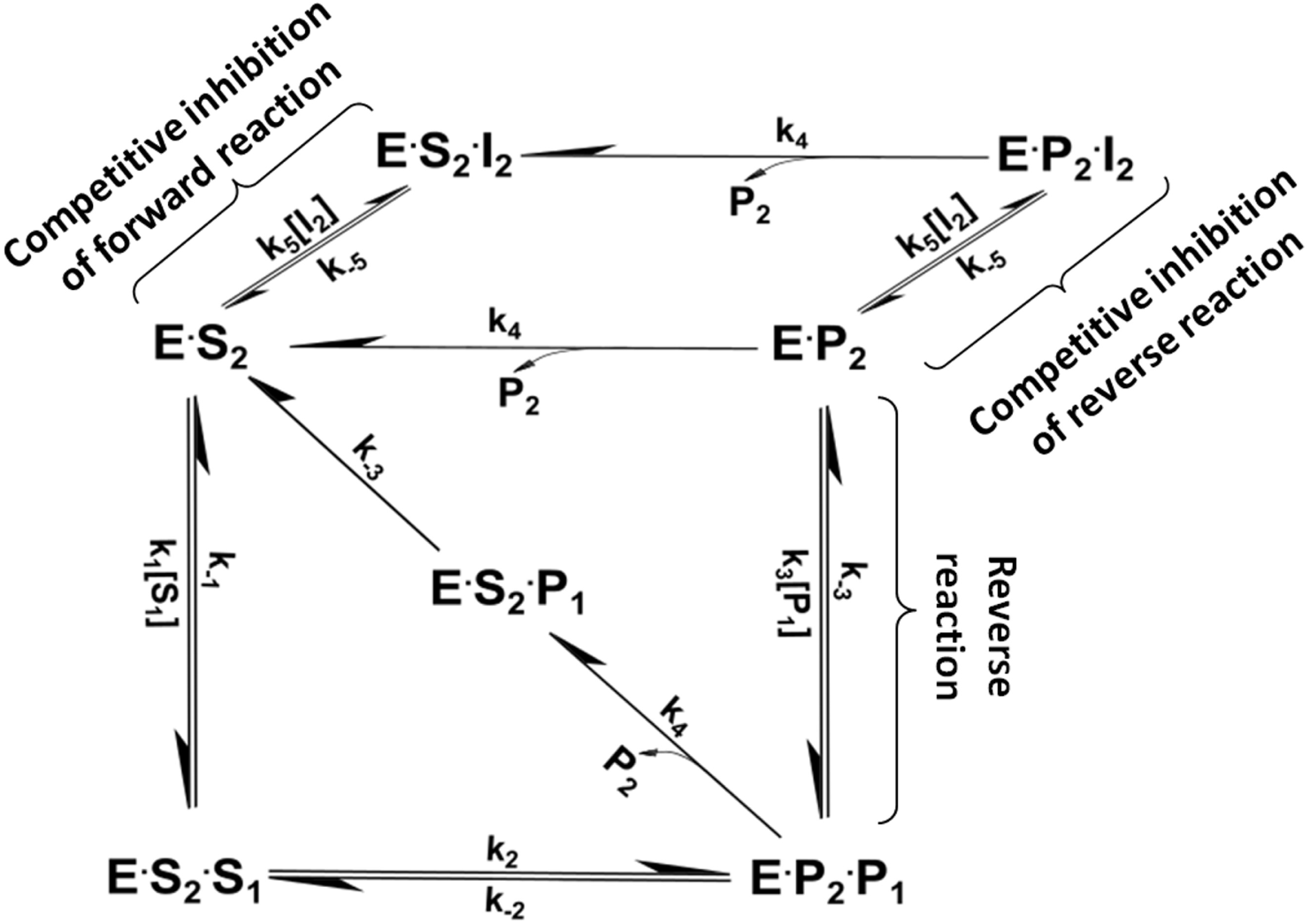
Reaction network for derepression of product inhibition in a two substrate two product enzyme

### 3.2 One substrate, one product enzymatic reactions

Similar principles can be applied to one substrate, one product enzymatic reactions. Fig. 2 displays the corresponding schematic for modulation of such reactions.

---

**Figure 2:**
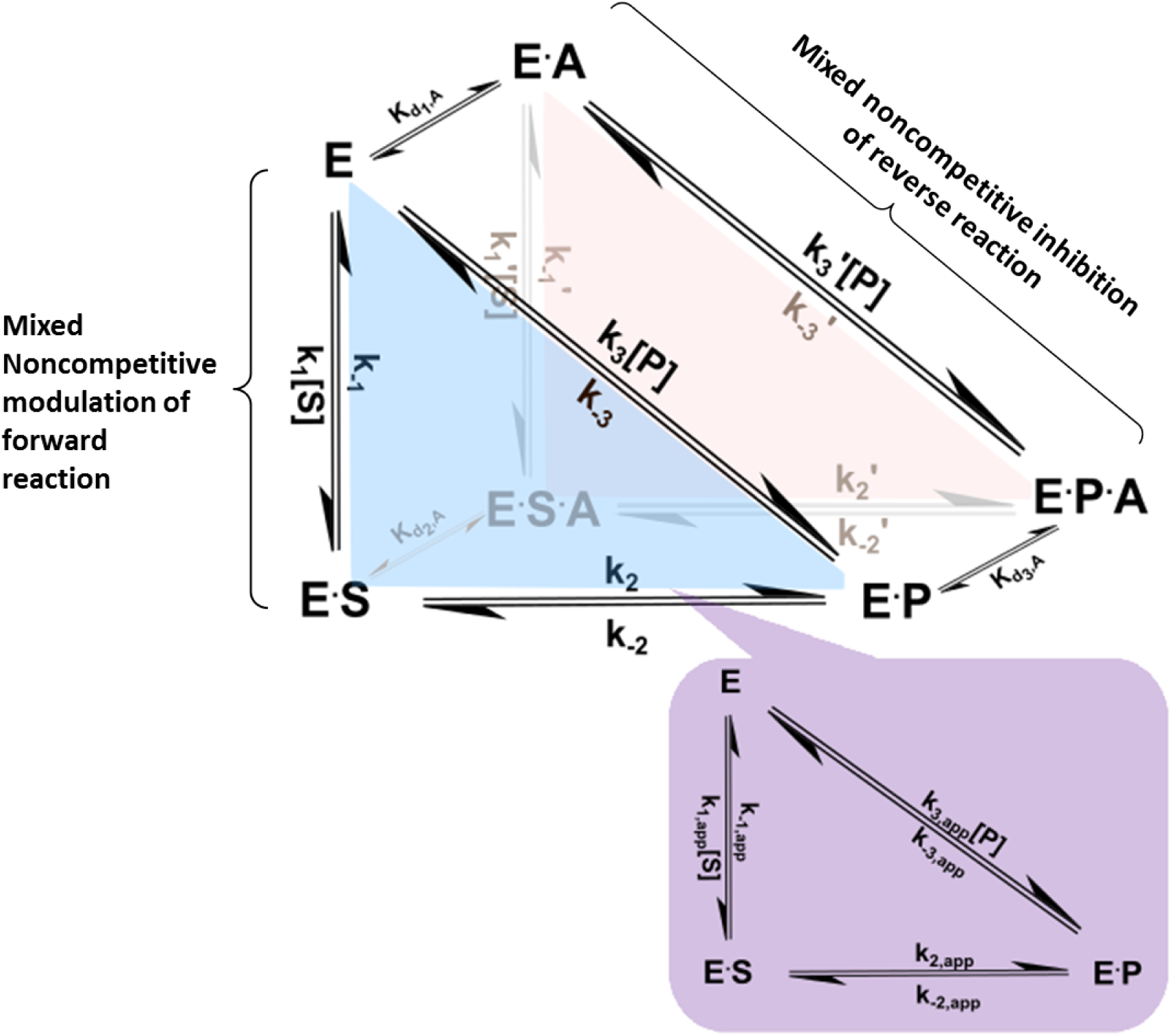
Reaction network for mechanism-based enzyme activation of a one substrate, one product reaction.

## Notes

### Competing Interest Statement

The authors have declared no competing interest.

